# A genome wide CRISPR screen reveals novel determinants of long-lived plasma cell secretory capacity

**DOI:** 10.1101/2025.02.28.640639

**Authors:** Lucas J. D’Souza, Jonathan N. Young, Heather Coffman, Edward P. Petrow, Deepta Bhattacharya

**Affiliations:** Department of Immunobiology, University of Arizona, Tucson, AZ; Department of Otolaryngology, University of Arizona, Tucson, AZ; Tucson Orthopedic Institute, Tucson, AZ

## Abstract

Plasma cell subsets vary in their lifespans and ability to sustain humoral immunity. We conducted a genome-wide CRISPR-Cas9 screen in myeloma cells for factors that promote surface expression of CD98, a marker of longevity in mouse plasma cells. A large fraction of genes found to promote CD98 expression in this screen are involved in secretory and other vesicles, including subunits of the V-type ATPase complex. Genetic ablation and chemical inhibition of V-type ATPases in myeloma cells and primary plasma cells, respectively, reduced antibody secretion. Mouse and human long-lived plasma cells had greater numbers of acidified vesicles than their short-lived counterparts, and this correlated with increased antibody secretory capacity. The screen also revealed a requirement for the signaling adapter MYD88 in CD98 expression. Plasma cell-specific deletion of *Myd88* led to reduced survival and antibody secretion by antigen-specific cells *in vivo* and unresponsiveness to BAFF and APRIL *ex vivo*. These data reveal novel regulators that link plasma cell secretory capacity and lifespan.

**Summary:** Long-lived plasma cells rely on V-type ATPases, PI4K, DDX3X, and MYD88 signals for maximal secretory capacity and survival

## Introduction

The humoral response following infection or vaccination involves recognition and activation of B cells by cognate antigen, followed by expansion, selection, and differentiation of these cells into quiescent memory B cells and antibody secreting plasma cells. If these antibodies are specific for protective epitopes on the pathogen, even small quantities can protect against otherwise lethal infections (Purtha et al., 2011). Thus, maintenance of protective antibody production is a key aspect of durable immunity. Yet, antibody production following some vaccines and infections are stable over time whereas others do so more transiently (Amanna et al., 2007; Olsson et al., 2007; Olotu et al., 2013; White et al., 2015). Underlying this variability is the lifespan of plasma cells that are generated.

Plasma cell subsets of varying longevity can be distinguished by using a combination of phenotypic markers. Human long-lived plasma cells tend to lack expression of CD19 (Mei et al., 2015; Halliley et al., 2015), though there are some exceptions to this rule (Brynjolfsson et al., 2017; Nguyen et al., 2024). In mice, a combination of markers such as BLIMP1 expression, B220, CD93, CXCR3, MHC-II, SLAMF6, LAG3, and the uptake of the fluorescent compound 2-deoxy-2-[(7-Nitro-2,1,3-benzoxadiazol-4-yl)-amino]-D-glucose (2NBDG) define plasma cell subsets of varying lifespans (D’Souza and Bhattacharya, 2019; Robinson et al., 2023; Lam et al., 2018; D’Souza et al., 2022). Another marker is the solute carrier protein CD98, which on the cell surface is subtly more highly expressed by long-lived plasma cells relative to their shorter-lived counterparts (Lam et al., 2018). CD98 is a heterodimer comprised of a heavy chain encoded by the *Slc3a2* gene and a light chain by the *Slc7a5* gene (Yanagida et al., 2001). This covalent pairing is not exclusive as the SLC3A2 protein can heterodimerize with other members of the SLC7 family to form a variety of transporters involved in amino acid uptake (Cantor and Ginsberg, 2012). Besides importing amino acids, CD98 can also interact with integrin β subunits to facilitate cell growth and survival (Feral et al., 2005). In plasma cells, the transcription factor BLIMP1 directly promotes *Slc7a5* expression (Tellier et al., 2016).

The activity of BLIMP1 is a major defining factor for inducing antibody secretory capacity in plasma cells. It does so in three distinct ways: first, by inducing a robust increase in transcription of *Igh* and *Igl* transcripts in plasma cells (Minnich et al., 2016); second, by upregulating the elongation factor ELL2 that permits alternative processing of the 3’ end of *Igh* transcripts and enabling translation of secretion-specific immunoglobulin heavy-chain proteins (Martincic et al., 2009); and lastly by inducing ATF4, ATF6, and XBP1 expression to support the high rate of antibody synthesis and folding as well as expansion of the ER network and secretory apparatus necessary for secretion (Tellier et al., 2016; Reimold et al., 2001; Shaffer et al., 2004; Hu et al., 2009; Taubenheim et al., 2012). Aside from BLIMP1, mTORC1 signaling is another key pathway that promotes antibody secretion (Jones et al., 2016; Benhamron et al., 2015; Brookens et al., 2020).

There is conflicting evidence of whether antibody secretory capacity and plasma cell lifespan are related or independent. On one hand, deletion of the transcription factor XBP-1 sharply reduces antibody secretory capacity but does not impact plasma cell differentiation or survival (Taubenheim et al., 2012). Similarly, genetic ablation of mTOR signaling in plasma cells reduces secretory capacity but not survival (Jones et al., 2016). These data demonstrate that there is no absolute requirement of antibody secretion for plasma cell survival. On the other hand, long-lived plasma cells tend to secrete more antibodies on a per cell basis than do their short-lived counterparts, but expression of XBP-1 and mTOR signaling are similar across these subsets (Lam et al., 2018). It therefore remains possible that some common pathways tune both lifespan and secretory capacity, perhaps explaining some of the functional heterogeneity in plasma cells. To identify such pathways, we performed an unbiased loss-of-function screen for genes that promote CD98 expression and investigated their roles in plasma cell function and longevity.

## Results

### CD98 expression on plasma cells correlates with 2NBDG uptake and is independent of secreted antibody isotype

We first characterized markers of plasma cell longevity that could potentially be used in genome-wide screens. In previous work, we showed that 2NBDG uptake correlated with the longevity of plasma cell subsets (Lam et al., 2018; D’Souza and Bhattacharya, 2019). Plasma cells that were 2NBDG+ also showed slightly higher levels of the large neutral amino acid transporter CD98, which is composed of SLC3A2 and SLC7A5, relative to their 2NBDG-counterparts. To test if immunoglobulin isotype usage impacts these correlations, we injected C57BL/6 (B6) mice with 2NBDG and enriched for CD138+ cells from spleens and bone marrow. A large fraction of splenic CD138+ B220+/-plasma cells were IgM+ while IgA+ plasma cells made up less than 10% of the pool **(Fig. S1 A)**. Consistent with previous reports and in contrast to the spleen (Wilmore et al., 2018), bone marrow plasma cells were mostly IgM+ or IgA+ **(Fig. S1 B)**. IgA+ plasma cells showed the highest frequency of 2NBDG+ cells followed by IgM+, and then IgG+ or IgE+ plasma cells in both spleen and bone marrow **(Fig. S1 C)**. When matched for antibody isotype, surface CD98 mean fluorescence intensity (MFI) was higher in plasma cells that took up 2NBDG (LLPCs) relative to plasma cells that were 2NBDG-(SLPCs) (**Fig. 1 A-C)**. Bone marrow LLPCs showed higher CD98 levels relative to their splenic counterparts. To quantify total, rather than just surface CD98 protein levels, we purified SLPCs and LLPCs from the spleens and bone marrows of B6 mice and examined for expression of CD98 following fixation and permeabilization. Again, bone marrow LLPCs had the highest total expression of CD98, followed by splenic LLPCs and then splenic SLPCs (**Fig. 1 D)**. Consistent with these findings, we used imaging flow cytometry to observe CD98 on both the surface and within the cytosol of fixed and permeabilized plasma cells in all groups (**Fig. 1 E)**. Bone marrow LLPCs showed the highest levels of CD98 on their surface as measured by similarity morphology indexes with CD138 as compared to both splenic plasma cell populations (**Fig. 1 E)**.

**Figure 1:**
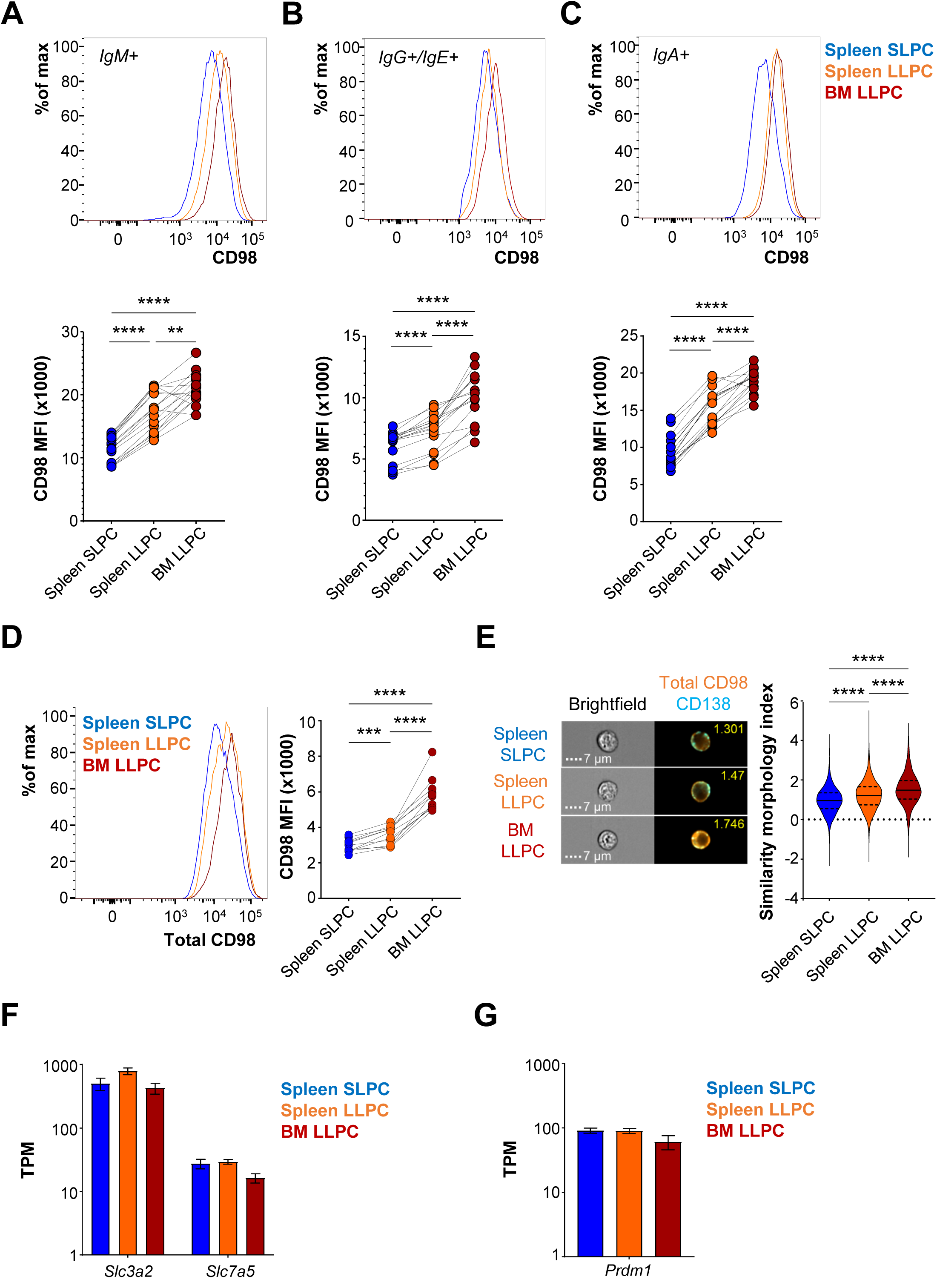
LLPCs have higher levels of CD98 than SLPCs despite having similar transcript reads. **(A-C)** Representative histograms (top) and quantification of mean fluorescence intensity (MFI) values (bottom) of CD98 expression on *ex vivo* (A) IgM+, (B) IgG+ and IgE+, and (C) IgA+ plasma cell subsets. Each data point represents cells from one mouse, and subsets from each mouse are connected by lines. Pooled data from 16 mice across 4 experiments. *p<0.05 by paired one-way ANOVA with Tukey’s correction. **(D)** Representative histogram of total (surface and internal) CD98 protein levels in sorted plasma cell subsets (left). Quantification of total CD98 MFI of each subset is represented (right). Each data point represents cells from one mouse, and subsets from each mouse are connected by lines. Pooled data from 11 mice across 3 experiments. *p<0.05 by paired one-way ANOVA with Tukey’s correction. **(E)** Cells in (D) were examined by imaging flow cytometry. Representative images for each subset are shown with surface CD138 (cyan) and total CD98 (orange) and their colocalization. Similarity morphology indices are indicated on the top right of the image and values for each subset are quantified. Representative graph of one of three independent experiments. *p<0.05 by unpaired one-way ANOVA with Games-Howell corrections. **(F)** Transcripts per million kilobase (TPM) values of CD98hc (*Slc3a2*) and CD98lc (*Slc7a5*) from Lam WY *et al* (Cell Reports 2018), showing values across splenic and bone marrow plasma cell subsets. Pooled data from three independent experiments. **(G)** Transcripts per million kilobase (TPM) values for BLIMP1 (*Prdm1*) from Lam WY *et al* (Cell Reports 2018) across splenic and bone marrow plasma cell subsets. Pooled data from three independent experiments.

Transcript abundances of *Slc3a2* and *Slc7a5*, as measured by RNA sequencing (Lam et al., 2018), were similar across plasma cell subsets (**Fig. 1 F)**. Transcript reads of *Prdm1*, which encodes for BLIMP-1, a known regulator of CD98 levels in plasma cells (Shi et al., 2015; Tellier et al., 2016), were also similar between short- and long-lived subsets (**Fig. 1 G)**. Thus, the differences between CD98 expression are apparent across plasma cell subsets irrespective of isotype and do not appear to be regulated at the level of transcription.

### A genetic screen to identify factors required for CD98 expression

In prior CRISPR approaches, we were unsuccessful in identifying genes that promote 2NBDG uptake (D’Souza et al., 2022). Given that elevated CD98 also marks long-lived plasma cell subsets, we carried out an unbiased genome wide CRISPR knockout screen for genes that promote its expression. We engineered the mouse myeloma cell line, 5TGM1, to express a Doxycycline inducible version of Cas9 (5TGM1-iCas9) **(Fig. S2 A)**. This strategy was chosen to allow for expansion of cells containing gRNAs that target essential genes which might otherwise have been lost in a constitutive Cas9 expression system. We then transduced 5TGM1-iCas9 cells with the Brie library containing 78,637 gRNAs targeting 19,674 genes of the mouse genome (Doench et al., 2016). After transduction, gRNAs were represented at levels similar to the input plasmid library (r^2^=0.57, **Fig. S2 B)**. We induced Cas9 expression by adding Doxycycline to the cell culture media and incubated cells for 7 days to ablate genes and promote complete depletion of residual proteins. From the Doxycycline treated cultures, we then sorted cells with reduced CD98 expression by FACS to purities >95% (**Fig. 2 A)**. Genomic DNA was then extracted from these cells and gRNA sequences amplified by PCR from the CD98-low and the total 5TGM1-iCas9-Brie populations. The resultant amplicons were sequenced and the abundance of gRNA was quantified relative to the total cell fraction using the Model-based Analysis of Genome-wide CRISPR/Cas9 Knockout (MaGeCK) algorithm (Li et al., 2014).

**Figure 2:**
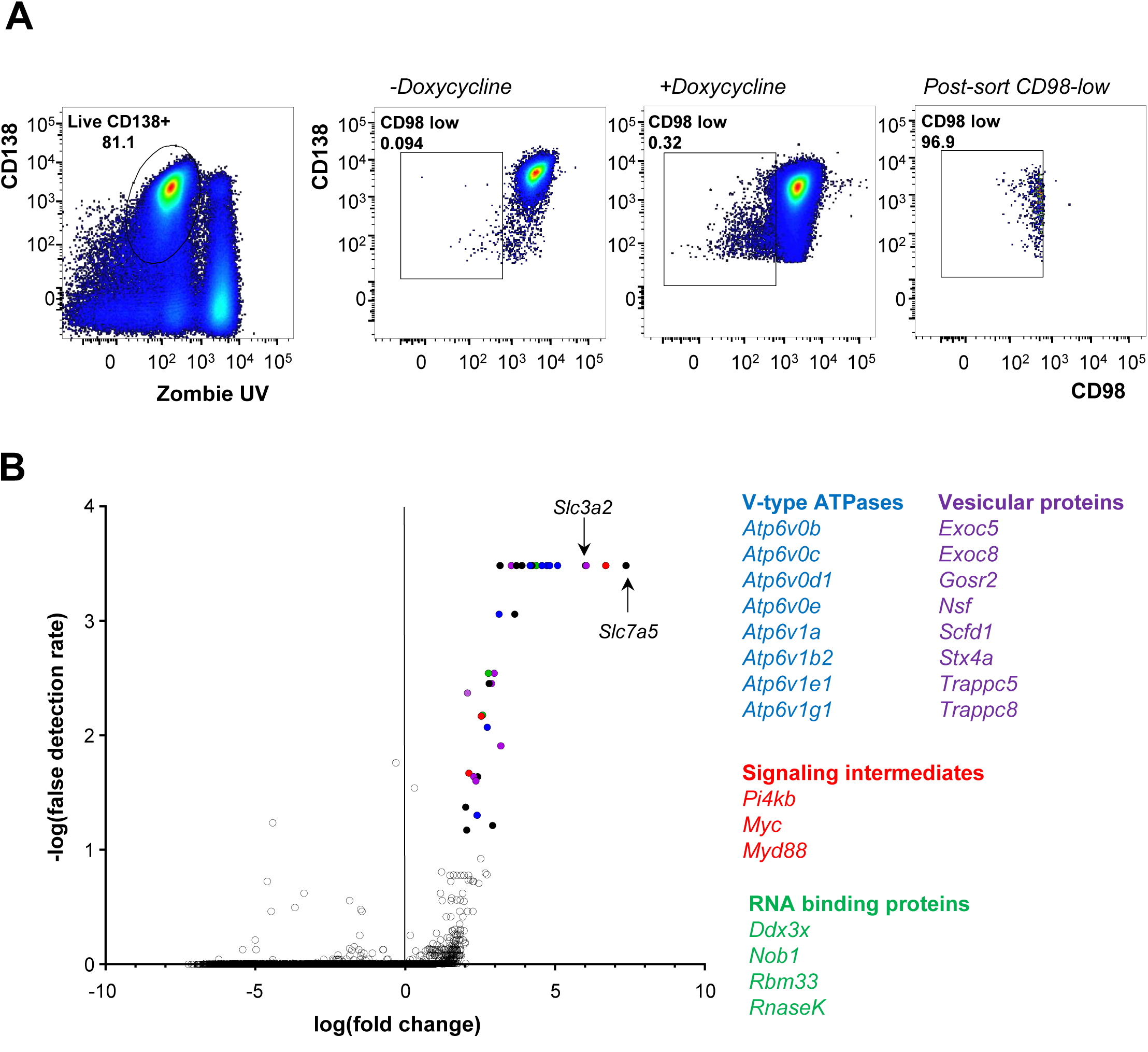
Genes involved in vesicle biology and exocytosis promote plasma cell CD98 expression. **(A)** Representative gating on 5TGM1-iCas9 transduced with the mouse Brie gRNA library. After treatment with Doxycycline for 7 days, 5TGM1-iCas9-Brie cells were gated first to exclude dead cells and debris (far left). On this gate, cells were subsequently gated as CD98-low based on the untreated 5TGM1-iCas9-Brie cells (center left). CD98-low cells in Doxycycline treated cultures (center right) were then purified by FACS to achieve a pure CD98-low fraction (far right). **(B)** MAGeCK analysis of genes enriched in the CD98-low fraction relative to unsorted 5TGM1-iCas9 cells transduced with the Brie library. Volcano plot of fold change between sorted cells against control cells are shown. Statistically significant genes are indicated as filled circles, the color of which corresponds to the class of genes indicated next to the volcano graph. Four biological replicates were analyzed.

In the CD98-low fraction, we observed 34 genes targeted by the gRNAs with greater than 2-fold enrichment and with a false discovery rate <0.05 (**Fig. 2 D)**. We observed enrichment of gRNAs targeting *Slc3a2* and *Slc7a5*, the genes that encode the heavy and light chains of the CD98 heterodimer, providing internal validation of the results of the genome-wide screen. Among the remaining significant 32 gRNA-targeted genes, we found 8 genes that associate with endosomal vesicles and 8 members of the vesicle-associated V-type ATPase complex. The remaining 16 genes include those that encode RNA export and processing proteins (*Thoc3*, *Ddx3x*, *Nob1*, *RnaseK*, and *Rbm33*), signaling intermediates (*Myd88*, *Myc*, and *Pi4kb*), proteins involved in genome structure (*Rad21* and *Smc3*), and the chaperone *Hsp90b1*. We did not identify *Prdm1* as a hit, despite its known role in promoting CD98 expression (Tellier et al., 2016). Yet when we compared cells treated with Doxycycline to untreated controls at day 7, we observed a drop in the read counts for gRNAs targeting *Irf4*, *Prdm1*, *Xbp1*, *Atf4*, and *Atf6b*, indicating a selection against cells that deleted these genes and potentially limiting their detection in our screen **(Fig. S2 C)**.

### Long-lived plasma cells have higher frequencies of acidic vesicles than do short-lived plasma cells

Given the enrichment of genes involved in multiple vesicular components and V-type ATPases in the screen, we chose to focus on these pathways first. The V-type ATPase is a multi-protein complex composed of a membrane bound proton pump (V0) powered by a cytosol-facing subunit (V1) that hydrolyzes ATP (Collins and Forgac, 2020). It acidifies lysosomes, endosomes, Golgi complexes, and secretory vesicles, all of which have significantly lower luminal pH relative to the cytosol (Casey et al., 2010). We purified mouse plasma cell subsets by FACS and quantified the frequencies of acidic vesicles using intracellular staining and imaging cytometry. Total ATP6V1A+ spot numbers per cell were slightly higher in bone marrow LLPCs relative to splenic LLPCs and SLPCs (**Fig. 3 A)**. To identify vesicular content differences more specifically between plasma cell subsets, we used characteristic markers. RAB7+ endosomes were present at higher frequencies in LLPCs of both spleen and bone marrow compared to SLPCs (**Fig. 3 B)**. Further, LAMP1+ lysosome frequencies per cell were highest in bone marrow LLPCs, followed by splenic LLPCs and SLPCs (**Fig. 3 C)**. We next examined if human plasma cell populations demonstrated similar trends for endosome and lysosome frequencies (gated as in **Fig. S3**). We observed higher frequencies of ATP6V1A+ and LAMP1+ vesicles in bone marrow plasma cells relative to tonsillar plasma cells (**Fig. 3 D-E)**. Put together, the frequencies of acidic vesicles in plasma cells are higher in subsets with longer lifespans.

**Figure 3:**
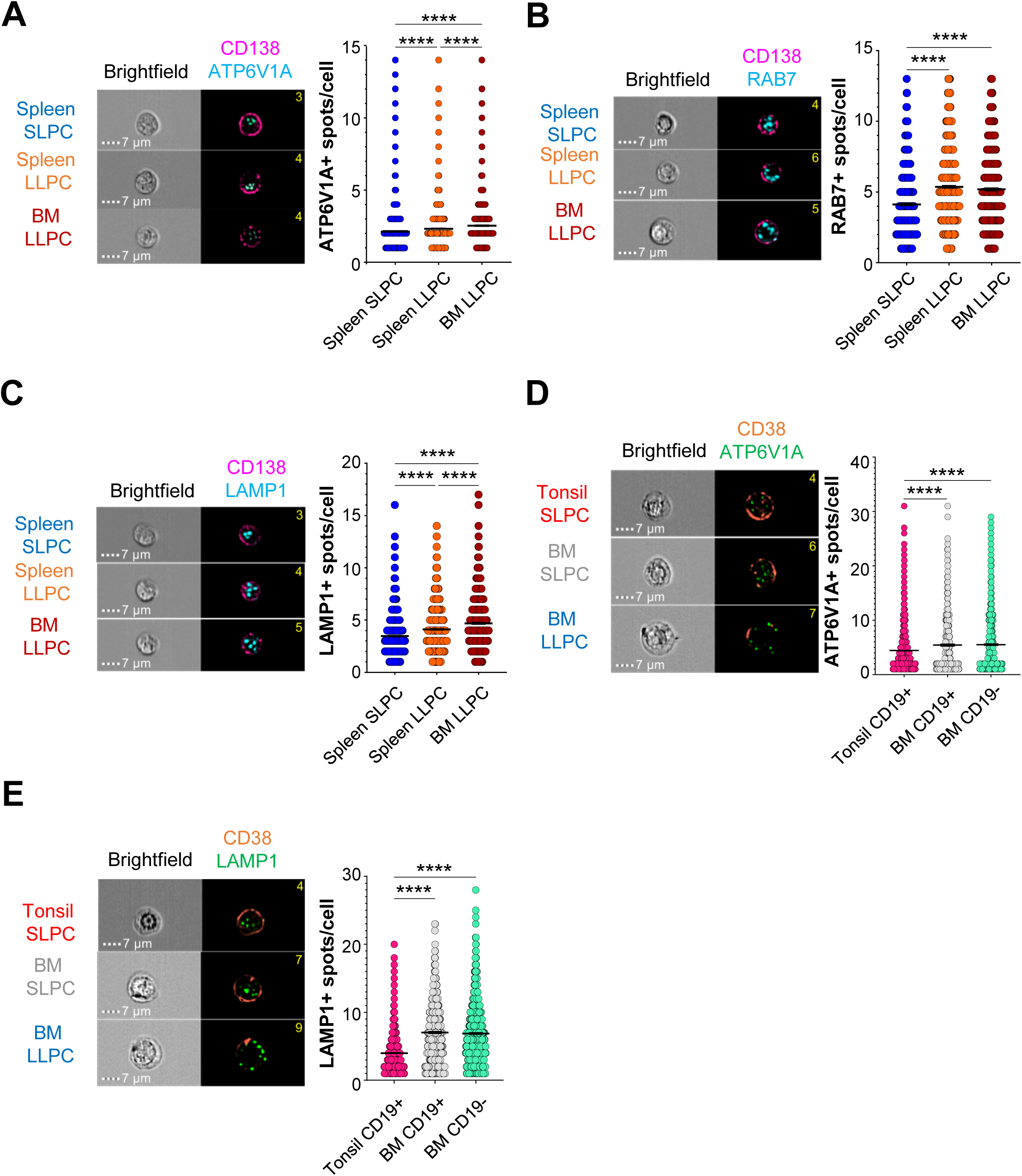
LLPCs have higher frequencies of acidic vacuoles than SLPCs. **(A)** Mouse plasma cell subsets from the spleen and bone marrow were sorted and stained for surface CD138 (pink) and intracellular ATP6V1A protein (cyan) and cells analyzer by imaging flow cytometry. Each circle represents one cell. ATP6V1A spot numbers/cell were enumerated and mean +/-SEM shown. Combined data from 9890-10755 cells from 12 mice across 3 experiments. *p<0.05 by unpaired one-way ANOVA with Games-Howell corrections. **(B)** Mouse plasma cell subsets as in (A) were stained for surface CD138 (pink) and RAB7 (cyan). RAB7+ spot numbers/cell were enumerated and mean +/-SEM graphed. Each circle represents one cell. Pooled data from 1639-2115 cells from 12 mice across 3 experiments. *p<0.05 by unpaired one-way ANOVA with Games-Howell corrections. **(C)** Mouse plasma cell subsets as in (A) were stained for surface CD138 (pink) and LAMP1 (cyan). LAMP1+ spot numbers/cell were enumerated and mean +/-SEM graphed. Each circle represents one cell. Pooled data from 3061-9159 cells from 12 mice across 3 experiments. *p<0.05 by unpaired one-way ANOVA with Games-Howell corrections. **(D)** Human plasma cells from the tonsil (CD19+ CD38+ CD27+ CD138-) and bone marrow CD19+ and CD19-plasma cells (CD27+ CD38+ CD138+) were stained for surface CD38 (orange) and intracellular ATP6V1A protein. ATP6V1A+ spot numbers/cell were quantified and mean +/-SEM plotted for all subsets. Each circle represents one cell. Combined data from 1156-9050 cells from 7 tonsil donors and 6 bone marrow donors across 3 experiments. *p<0.05 by unpaired one-way ANOVA with Games-Howell corrections. **(E)** Human plasma cell subsets as in (D) were stained for surface CD38 and intracellular LAMP1. LAMP1+ spot numbers/cell were quantified and mean +/-SEM plotted. Each circle represents one cell. Pooled data from 2614-10742 cells from 9 tonsil donors and 7 bone marrow donors from 4 experiments. *p<0.05 by unpaired one-way ANOVA with Games-Howell corrections.

We reasoned that these differences in vesicular content might reflect differential antibody secretory capacities of plasma cell subsets. We sorted polyclonal SLPCs and LLPCs from the spleens and bone marrows of mice and seeded equal numbers for ELISpot assays, as spot size can serve as a surrogate of antibody secretory capacity (Jones et al., 2016). We found that bone marrow LLPCs yielded spots with the largest average area as compared to their splenic equivalents irrespective of antibody isotype (**Fig. 4 A-D)**. Splenic SLPC spots were the smallest in comparison. Similarly, irrespective of isotype, we found that human CD19-bone marrow plasma cells secreted more antibodies per cell than did CD19+ bone marrow plasma cells (**Fig. 4 E-H)**, which on an average have shorter lifespans than their CD19-counterparts. These findings suggest that for both mice and humans and across isotypes, long-lived plasma cells have higher numbers of acidic vesicles and secrete more antibodies than their shorter-lived counterparts.

**Figure 4:**
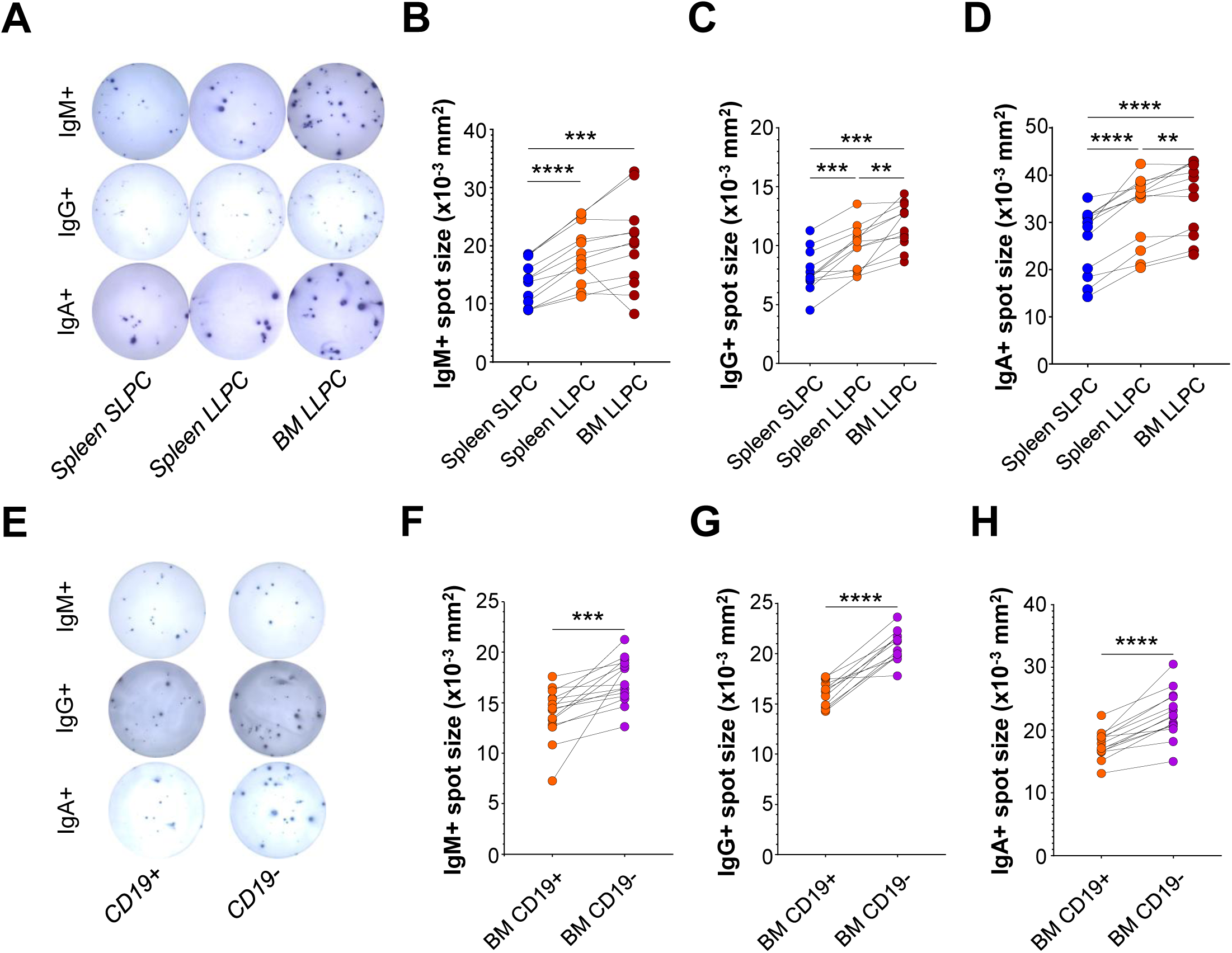
LLPCs secrete more antibodies than SLPCs independent of immunoglobulin isotype. **(A-D)** IgM+ *ex vivo* splenic SLPCs, LLPCs, and bone marrow LLPCs were sorted and seeded into wells of an ELISpot plate. (A) Representative images showing spots for each subset and isotype are indicated. Quantification of **(B)** IgM+, **(C)** IgG+, and **(D)** IgA+ plasma cell spot area of each subset is represented. Each data point represents mean spot sizes from one mouse, and subsets within each mouse are connected by lines. Pooled data from 12 mice across 3 experiments. *p<0.05 by paired one-way ANOVA with Tukey’s correction. **(E-H)** Human bone marrow plasma cells (CD27+ CD38+ CD138+) were sorted as CD19+ and CD19-populations and seeded into wells of an ELISpot plate. (E) Representative images showing spots for CD19+/-plasma cells secreting the indicated isotype are shown. Quantification of spot area of (F) IgM+, (G) IgG+, and (H) IgA+ plasma cells are represented. Each data point represents one donor, and subsets from an individual are connected by lines. Pooled data from 15 donors across 4 experiments for IgM, 12 donors across 3 experiments for IgG, and 14 donors across 4 experiments for IgA. *p<0.05 by paired t tests.

### Ablation of V-type ATPase function reduces antibody secretion

We next addressed if there were a direct link between vesicular pH and the antibody secretion capacity of plasma cells. In the first approach, we chemically inhibited V-type ATPase activity using Bafilomycin A1 (Bowman et al., 1988). To detect changes in vesicular pH in cells, we transduced Cas9-expressing 5TGM1 cells (5TGM1-Cas9) with the pH Lysosomal Activity Reporter (pHLARE) construct (Webb et al., 2021). The resultant 5TGM1-Cas9-pHLARE cells express *R. norvegicus* LAMP1 as a fusion protein with the pH-sensitive superfolder GFP (sfGFP) in the lumen of the lysosome and the pH-insensitive mCherry on the cytosol side of the lysosome, allowing for ratiometric calculation of lysosomal pH (Pédelacq et al., 2006). We observed a 1.3- to 5.6-fold increase in the ratio of sfGFP to mCherry in these cells with increasing concentrations of Bafilomycin A1 (**Fig. 5 A)**. Reciprocally, the MFI of surface CD98 was proportionally reduced in all groups treated with Bafilomycin A1 (**Fig. 5 B)**. Culture supernatants from groups treated with higher doses of the inhibitor showed reduced antibody levels relative to untreated controls (**Fig. 5 C)**. Lastly, we treated purified plasma cell subsets with this inhibitor and observed a reduction in the spot size of secreted immunoglobulin per cell (**Fig. 5 D)**. These findings indicate that pharmacological inhibition of V-type ATPase activity impairs both surface CD98 expression and antibody secretion.

**Figure 5:**
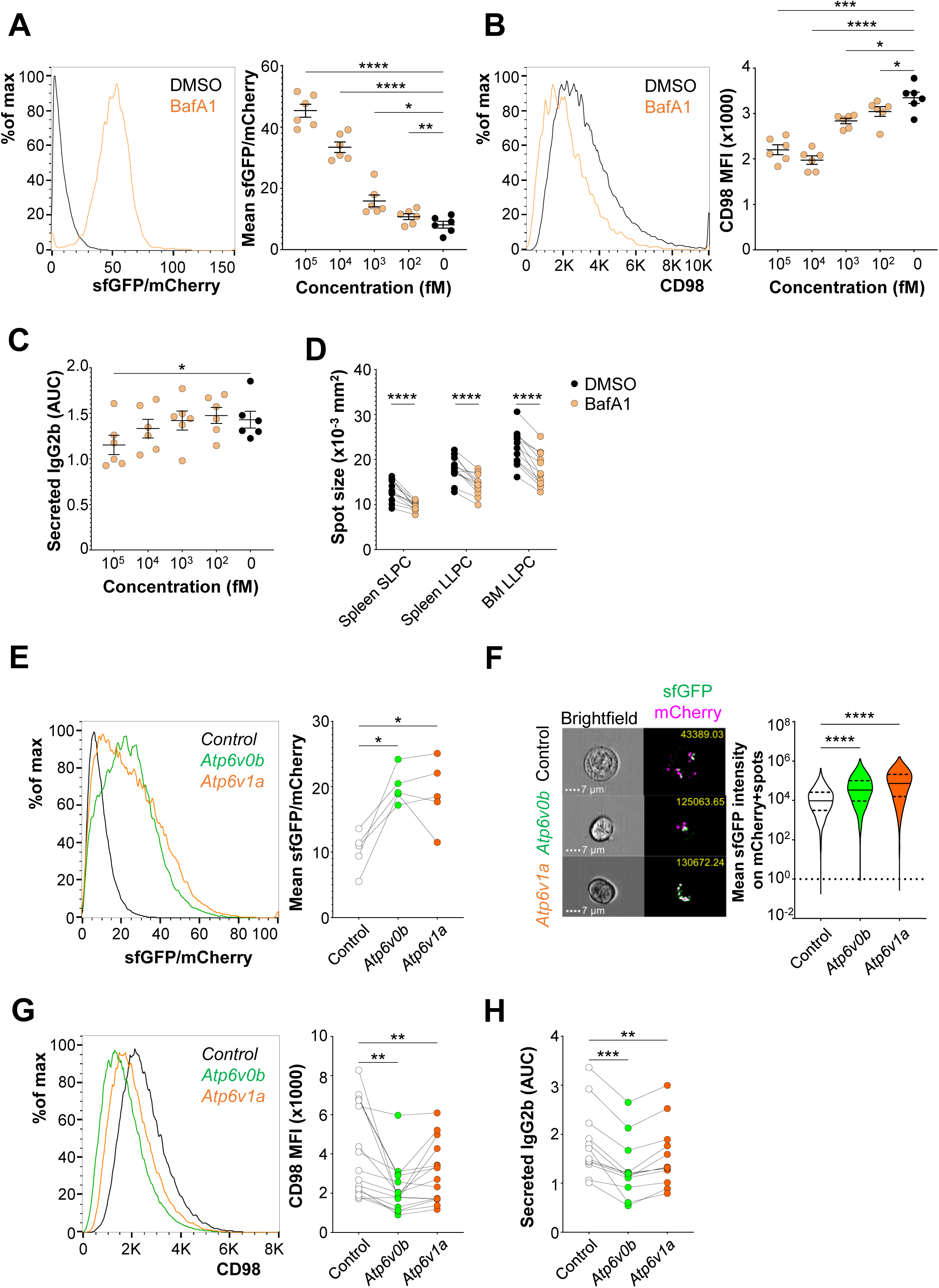
V-type ATPase activity promotes CD98 expression and antibody secretion. **(A)** 5TGM1-Cas9-pHLARE cells were treated overnight with the indicated doses of Bafilomycin A1. Representative histogram of the ratio of sfGFP to mCherry MFI in DMSO (black) and 100nM Bafilomycin A1 (tan) treated groups (left). Quantification of mean sfGFP/mCherry ratio (right) in control and knockout cultures is shown. Each circle represents the MFI of cells in one well. Pooled data from 6 experiments. *p<0.05 by repeated measures one-way ANOVA with Dunnett’s correction. **(B)** Cells in (A) were examined for MFI of surface CD98 by flow cytometry. Each circle represents the mean values for cells from a single well of an experiment. Pooled data from 6 experiments. *p<0.05 by repeated measures one-way ANOVA with Dunnett’s correction. **(C)** Culture supernatants of cells in (A) were scored for secreted IgG2b by ELISA. AUC for each group is shown. Pooled data from 6 independent experiments. *p<0.05 by repeated measures one-way ANOVA with Dunnett’s correction. **(D)** *Ex vivo* plasma cell subsets were sorted, and equal numbers were seeded into wells of an ELISpot plate containing media (untreated, black) or media with 10nM Bafilomycin A1 (tan). Quantification of average spot area of each subset in each treatment is shown. Each circle represents data from a single mouse and is connected across treatments with a line. Pooled data from 12 mice from 3 experiments. *p<0.05 by two-way ANOVA. **(E)** 5TGM1-Cas9-pHLARE cells were transduced with lentivirus containing gRNA targeting *Atp6v0b* or *Atpv1a* along with a control gRNA. Representative histogram (left) of the ratio of sfGFP to mCherry MFI in control gRNA (black), *Atp6v0b*-(green), and *Atp6v1a*-targeting gRNA (orange) groups. Quantification of mean sfGFP/mCherry ratio (right) in control and knockout cultures. Each circle represents a single treatment and data points are joined by a line within the same experiment. Pooled data from 5 independent experiments. *p<0.05 by paired one-way ANOVA with Tukey’s correction. **(F)** Groups in (E) were assessed for sfGFP intensity on mCherry spots per cell by imaging flow cytometry. Representative images of each group (left) at 60x are shown with ratio values in the top right. Mean ratio of GFP intensity on mCherry+ spots/cell were quantified. Each circle represents one cell from untreated/treated groups. Representative graph of 3 independent experiments. *p<0.05 by unpaired one-way ANOVA with Games-Howell corrections. **(G)** CD98 expression on 5TGM1-Cas9 cells transduced with control gRNA (black), *Atp6v0b*-(green), and *Atp6v1a*-targeting gRNA (orange). Representative histogram (left) and quantified CD98 MFI (right) depicted. Each circle represents the MFI of cells in one well and groups from the same experiment are connected by a line. Combined data from 15 experiments. *p<0.05 by paired one-way ANOVA with Tukey’s correction. **(H)** Secreted IgG2b from culture supernatant of groups in (G) was quantified by ELISA. Area under the curve (AUC) for each group is shown. Pooled data from 12 experiments. *p<0.05 by paired one-way ANOVA with Tukey’s correction.

We confirmed these findings genetically by ablating V-type ATPase activity in 5TGM1 cells using CRISPR-Cas9. We targeted *Atp6v0b* and *Atp6v1a* as representative genes of the V0 and V1 subunit of the ATPase complex. Targeting these genes led to a nearly two-fold increase in the ratio of sfGFP to mCherry, indicating a rise in lysosomal pH (**Fig. 5 E)**. This increase in sfGFP fluorescence can be traced to increased sfGFP intensity on mCherry+ spots in cells as observed by imaging cytometry (**Fig. 5 F)**. In both cases, we observed a drop in surface CD98 levels as well as secreted antibodies in their culture supernatants (**Fig. 5 G-H)**. Further, the survival of *Atp6v0b*- and *Atp6v1a*-deficient myeloma cells in culture was compromised relative to control cells **(Fig. S4)**. In conclusion, V-type ATPases promote CD98 expression, survival, and antibody secretory capacity of plasma cells.

### Pi4kb promotes vesicular acidification and antibody secretion in plasma cells

Among the genes enriched in the CD98-low fraction was *Pi4kb*, the gene that encodes the enzyme phosphatidylinositol-4-hydroxide kinase B (PI4KB). It is a type-III PI4K that catalyzes the formation of phosphoinositide-4-phosphate (PI4P) at Golgi surfaces which subsequently promotes vesicle formation (Meyers and Cantley, 1997; Godi et al., 1999; Baba and Balla, 2020). We ablated *Pi4kb* in 5TGM1-Cas9 cells using CRISPR-Cas9 and observed a drop in the total numbers of ATP6V1A+ spots per cell and LAMP1+ lysosomes per cell (**Fig. 6 A-B)**. Deletion of *Pi4kb* in 5TGM1-Cas9-pHLARE cells showed lysosomes with higher sfGFP signal (**Fig. 6 C)**, suggesting that loss of PI4KB leads to poor maintenance of V-type ATPase complexes and vesicular acidification. *Pi4kb*-deficient 5TGM1 cells also showed reduced surface CD98 expression, secreted fewer antibodies, and had poor survival in culture (**Fig. 6 D-E and Fig. S4)**. Thus, PI4KB promotes the formation and acidification of intracellular vesicles and antibody secretory capacity through V-type ATPase function.

**Figure 6:**
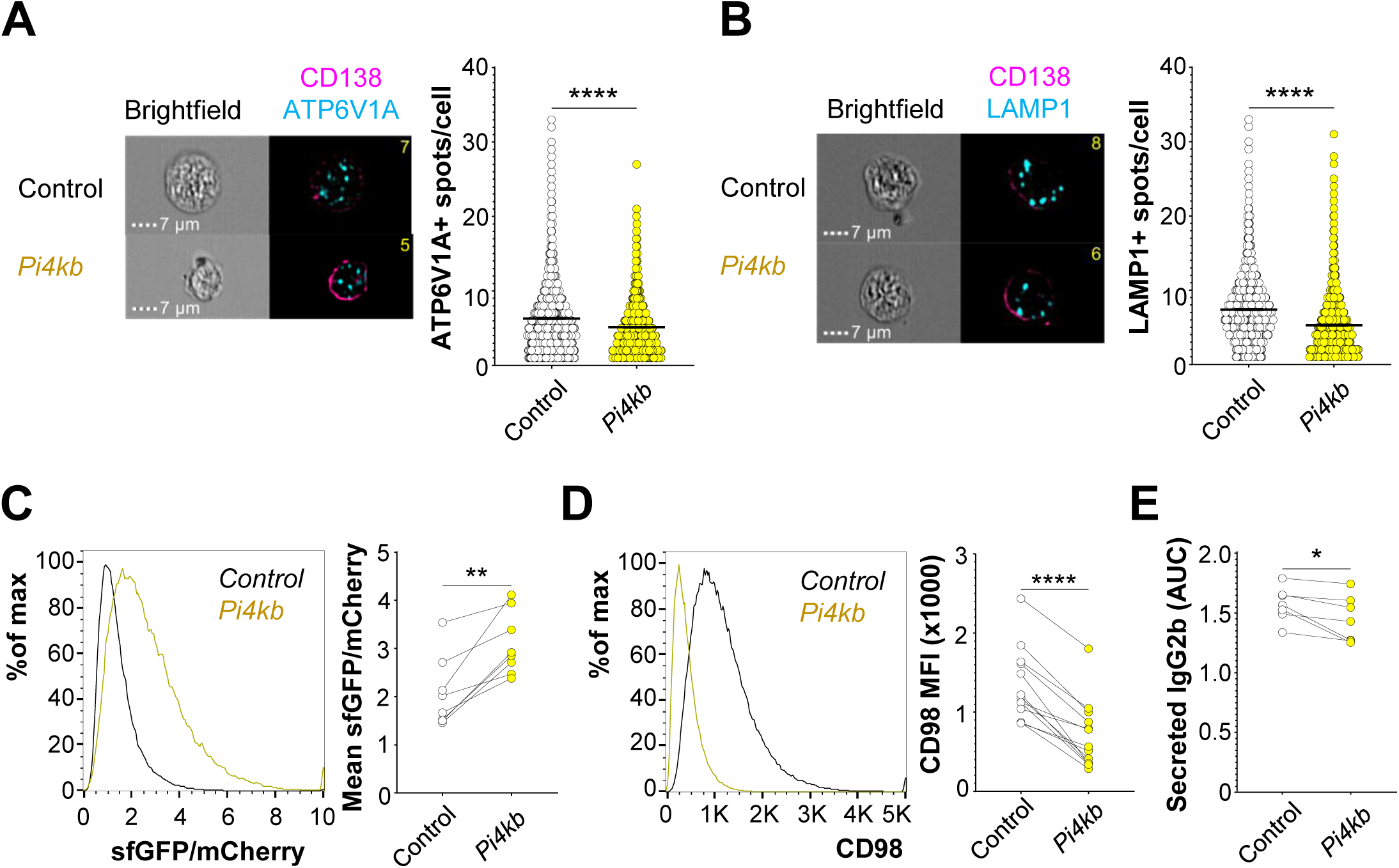
Ablation of *Pi4kb* reduces acidic vacuoles and antibody secretion. **(A)** 5TGM1-Cas9 cells transduced with control gRNA (black) and *Pi4kb*-targeting gRNA (yellow) and then scored for intracellular ATP6V1A+ spots/cell. Representative images for each of the groups (left) and numbers of ATP6V1A+ spots/cell (right) were quantified. Each circle represents one cell. Pooled data from 4236-4368 cells across 3 independent experiments. *p<0.05 by an unpaired t-test with Welch’s correction. **(B)** Groups in (A) were scored for intracellular LAMP1+ spots/cell. Representative images for each of the groups (left) and numbers of LAMP1+ spots/cell (right) were quantified. Each circle represents one cell. Pooled data from 4830-4837 cells across 3 independent experiments. *p<0.05 by an unpaired t-test with Welch’s correction. **(C)** 5TGM1-Cas9-pHLARE cells were transduced with lentivirus containing gRNA targeting *Pi4kb* along with a control gRNA. Representative histogram (left) of the ratio of sfGFP to mCherry MFI in control gRNA (black) and *Pi4kb*-targeting gRNA (yellow) groups. Quantification of mean sfGFP/mCherry ratio (right) in control and knockout cultures. Each circle represents values from a single group and groups within the same experiment are joined by a line. Pooled data from 7 independent experiments. *p<0.05 by a paired t-test. **(D)** Surface CD98 expression on groups in (C). Representative histogram (left) and quantified CD98 MFI (right) are shown. Combined data from 13 experiments. *p<0.05 by a paired t-test. **(E)** Secreted IgG2b from culture supernatant of groups in (C) were quantified by ELISA. AUC for each group is shown. Pooled data from 7 experiments. *p<0.05 by a paired t-test.

### DDX3X affects antibody secretory capacity of plasma cells independent of vesicular acidification

The ATP-dependent DEAD-box RNA helicase *Ddx3x* was another highly enriched gene in the CD98-low fraction (**Fig. 2 B)**. The DDX3X protein associates with complex secondary structures in mRNA and enables protein synthesis (Lai et al., 2008; Soto-Rifo et al., 2012; Calviello et al., 2021). Deletion of *Ddx3x* in 5TGM1-Cas9 cells reduced surface CD98 levels and decreased secreted immunoglobulin in culture supernatants (**Fig. 7 A-B)**. Ablating *Ddx3x* expression in 5TGM1-Cas9-pHLARE cells did not affect lysosomal pH levels, suggesting that it regulates the antibody secretory capacity of cells through a mechanism distinct from the PI4KB-V-type ATPase pathway (**Fig. 7 C)**.

**Figure 7:**
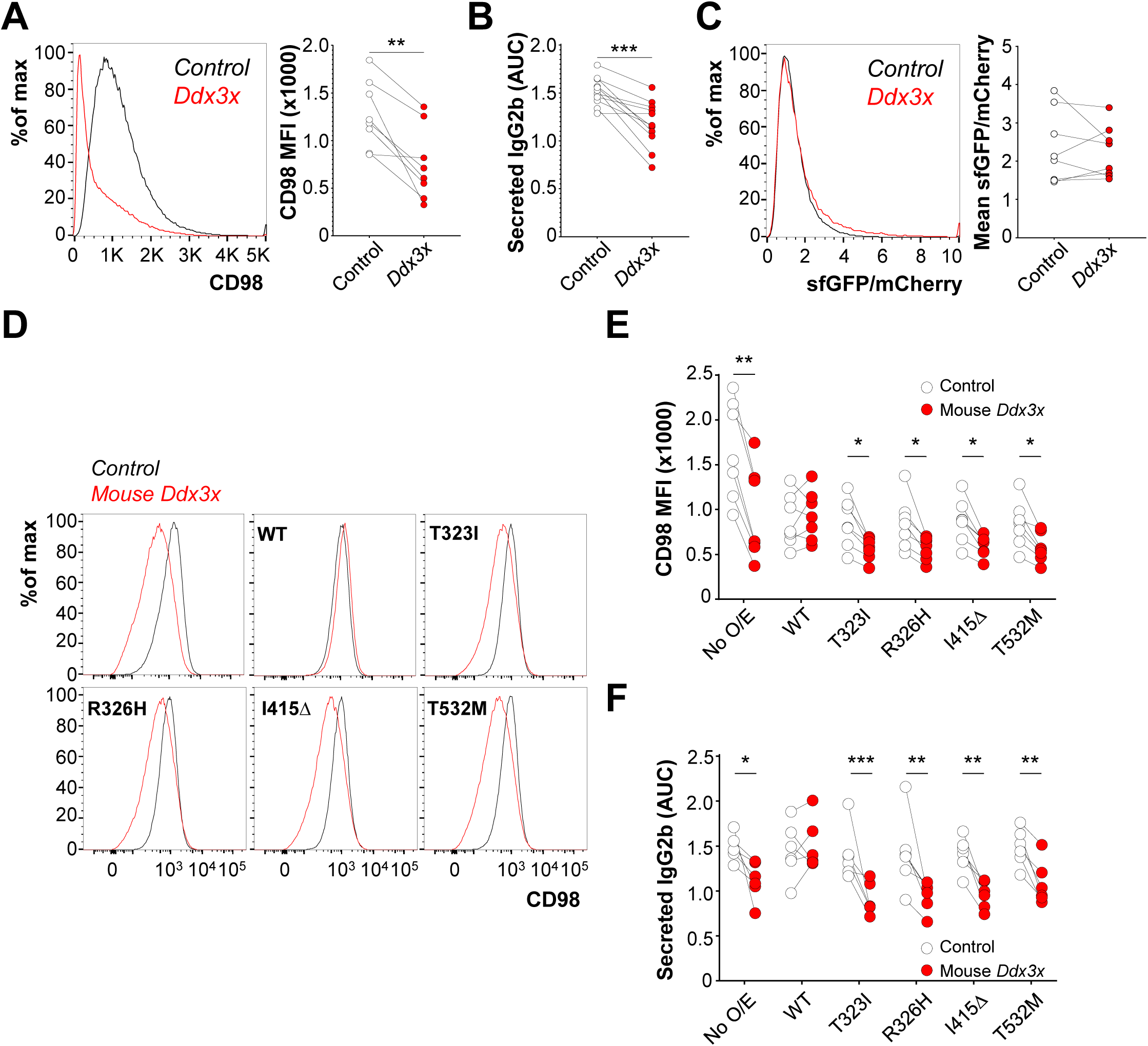
DDX3X regulates antibody secretion independent of vacuolar acidification. **(A)** Surface CD98 expression on 5TGM1-Cas9 cells transduced with control gRNA or gRNA targeting *Ddx3x*. Representative histogram (left) and quantified CD98 MFI (right) are shown. Each circle represents values from cells in one well and are connected with groups from the same experiment with a line. Combined data from 8 experiments. *p<0.05 by a paired t-test. **(B)** IgG2b antibody levels in culture supernatants of cells in (A). AUC for each group is shown. Pooled data from 11 independent experiments. *p<0.05 by a paired t-test. **(C)** 5TGM1-Cas9-pHLARE cells were transduced with lentivirus containing gRNA targeting *Ddx3x* as well as a control gRNA. Representative histogram (left) of the ratio of sfGFP to mCherry MFI in control gRNA (black) and *Ddx3x*-targeting gRNA (red) groups. Quantification of mean sfGFP/mCherry ratio (right) in control and knockout cultures is shown. Each circle represents a single group and groups within the same experiment are joined by a line. Pooled data from 8 experiments. No significance observed by a paired t-test. **(D)** 5TGM1-Cas9 cells overexpressing hsDDX3X or catalytically inactive hsDDX3X mutants were transduced with control gRNA or gRNA targeting mouse *Ddx3x*. Representative histograms for surface CD98 distribution on cells transduced with control gRNA (black) and mouse *Ddx3x* (red) are shown for cells without overexpression (top left), cells overexpressing WT DDX3X (top center), and cells overexpressing indicated mutant DDX3X (remaining squares). **(E)** Quantification of surface CD98 expression on cells in (D) as well as cells overexpressing DDX3X mutants. Each circle represents a single group and groups within the same experiment are joined by a line. Pooled data from 6 independent experiments. *p<0.05 by two-way ANOVA with Sidak’s correction. **(F)** Culture supernatants of all groups in (D) were assessed for secreted IgG2b by ELISA. AUC for each group is shown. Pooled data from 6 experiments. *p<0.05 by two-way ANOVA with Sidak’s correction.

*Ddx3x*-deficient myeloma cells also survived poorly in culture over time **(Fig. S4)**. We next lentivirally transduced 5TGM1-Cas9 cells with wild-type human DDX3X (hsDDX3X) or versions with point mutations in their helicase domain (T323I, R326H) or aberrations in their ATP-binding domain (I415Δ, T532M) that have previously been reported in patients with DDX3X syndrome (Lennox et al., 2020). We then depleted murine wild-type DDX3X in these cells using a gRNA that targeting mouse *Ddx3x* but not the transgenic hs*Ddx3x* cassette. At 5 days post-transduction, we found that deleted cultures were rescued by unmodified hsDDX3X (WT) as measured by surface CD98 levels (**Fig. 7 D-E)**. When quantified by ELISA, these cultures also showed similar levels of secreted immunoglobulin in their supernatants relative to untargeted control cells, indicating that expression of hsDDX3X overcame the defect otherwise seen with ablation of endogenous DDX3X (**Fig. 7 F)**. In all cases, overexpression of hsDDX3X mutants failed to rescue surface CD98 levels or secreted immunoglobulin in culture supernatants (**Fig. 7 D-F)**. This observation is not because of differential levels of DDX3X overexpression, as they were found to be comparable across all groups when probed for intracellular DDX3X protein **(Fig. S5)**. Put together, both the ATPase and helicase domain of DDX3X are necessary for proper antibody secretion in plasma cells.

### MYD88 is a plasma cell intrinsic determinant of antibody secretion capacity

We observed an enrichment of gRNAs targeting the signaling intermediate MYD88 in the CD98-low fraction (**Fig. 2 B)**. MYD88 is an adaptor protein required for transducing signals through all Toll-like receptors (TLRs), the IL-1 receptor (IL-1R), and cytokine signals through Transmembrane Activator and CAMI Interactor (TACI) by TRAF6-dependent and -independent mechanisms (Takeda and Akira, 2004). As *Myd88*-germline knockout (KO) mice are viable and do not show any defects in B cell development (Pasare and Medzhitov, 2005), we first examined plasma cell subsets for CD98 expression. We injected *Myd88*-KO and age- and sex-matched C57BL/6N mice with 2NBDG and examined plasma cell subsets from the spleens and bone marrows of these mice. Across all 2NBDG-subsets, *Myd88*-KO plasma cells had lower expression of surface CD98 (**Fig. 8 A)**. We then seeded equal numbers of plasma cell subsets into an ELISpot assay. As signals downstream of MYD88 promote class-switch recombination (He et al., 2010), we compared only IgM-expressing plasma cell subsets. IgM+ spot areas were reduced only in splenic and bone marrow LLPCs, suggesting that MYD88 signals promote antibody secretion in longer-lived plasma cell subsets (**Fig. 8 B)**.

**Figure 8:**
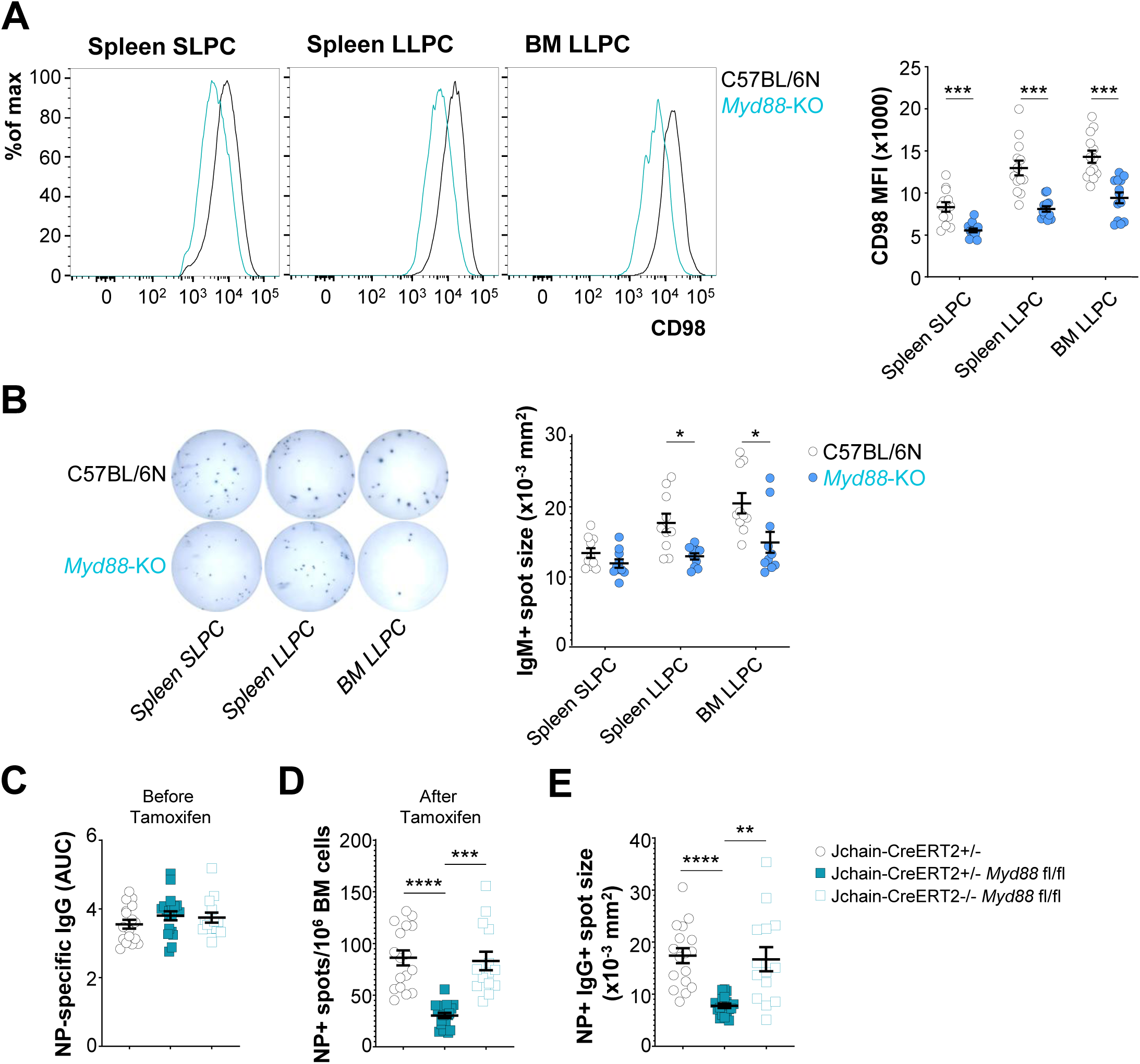
Plasma cell intrinsic MYD88 promotes antibody secretory capacity and longevity. **(A)** Representative histograms of surface CD98 levels on IgM+ splenic SLPCs (left), splenic LLPCs (center), and bone marrow LLPCs (right) from C57BL/6N (black) and *Myd88*-KO mice (blue). CD98 MFI was then quantified on indicated groups. Each circle is representative of a single mouse. Pooled data from 13 mice of each genotype across 4 independent experiments. *p<0.05 by two-way ANOVA with Sidak’s correction. **(B)** Indicated IgM+ plasma cell populations were purified from the spleens and bone marrows of C57BL/6N (black) and *Myd88*-KO mice (blue) and seeded into an ELISpot plate. Representative images showing spots from indicated plasma cell subset from indicated mice are shown (left). Spot sizes of IgM-secreting plasma cells are quantified (right), where each circle represents plasma cells from a single mouse. Combined data from 10 mice across 3 experiments. *p<0.05 by two-way ANOVA with Sidak’s correction. **(C)** Serum NP-specific IgG levels in Jchain-CreERT2+/-(black, filled circles), Jchain-CreERT2+/-*Myd88* fl/fl (blue, filled squares), and Jchain-CreERT2-/-*Myd88* fl/fl (blue, open squares) at 8 weeks post immunization with NP-CGG in Alhydrogel. Each square represents a single mouse. Pooled data from 14-20 mice per group across 3 different experiments. No statistical significance observed by repeated measures one-way ANOVA with Dunnett’s correction. **(D)** Mice in (C) were placed on tamoxifen chow for 14 days. CD138+ bone marrow cells were enriched and seeded into an ELISpot plate. Numbers of NP-specific IgG+ spots per million cells are quantified for each mouse and represented. *p<0.05 by repeated measures one-way ANOVA with Dunnett’s correction. **(E)** NP-specific IgG spot sizes in (D) were quantified and shown. *p<0.05 by repeated measures one-way ANOVA with Dunnett’s correction.

To test if MYD88 affects antibody secretory capacity in a plasma cell intrinsic manner, we generated lineage specific *Myd88* knockout mice. We bred mice with loxP sites flanking exon 3 of the *Myd88* genomic locus (*Myd88* fl/fl) with mice carrying an IRES-CreERT2 cassette at the 3’ UTR of the *IgJ* locus (Jchain-CreERT2). In these mice, treatment with tamoxifen induces deletion of *Myd88* in plasma cells without impacting other cell types (Hou et al., 2008; Wong et al., 2020). We immunized these mice (Jchain-CreERT2+/-*Myd88* fl/fl) along with littermate Cre-negative (Jchain-CreERT2-/-*Myd88* fl/fl) and Cre-only (Jchain-CreERT2+/-) control mice with 100µg NP-CGG in 1% Alhydrogel, an adjuvant that does not activate TLR signals (Gavin et al., 2006). At 8 weeks, we observed comparable levels of serum NP-specific antibodies across all groups (**Fig. 8 C)**. We then fed mice Tamoxifen chow over the course of 2 weeks to induce deletion of *Myd88*. Bone marrow LLPCs from these treated mice were then enriched and examined for spot frequencies and sizes in ELISpot assays. We observed a significant reduction in both the numbers and spot sizes of NP-specific plasma cells in the bone marrows of Jchain-CreERT2+/-*Myd88* fl/fl mice relative to both control mouse groups (**Fig. 8 D-E)**. This suggests that plasma cell intrinsic MYD88 signaling promotes both survival and secretory capacity of plasma cells *in vivo*.

### Cytokine signals through TACI calibrate antibody secretory capacity in a MYD88 fashion

To identify which MYD88-dependent signals promote antibody secretion, we examined a previously generated RNAseq databases for expression of receptors that signal through MYD88 in plasma cells and myeloma cells (Lam et al., 2018; D’Souza et al., 2022). We observed no detectable levels of *Il1r* transcript in all populations examined, but high expression of *Tnfrsf13b*, the gene that encodes TACI, in all groups **(Fig. S6 A)**. Further, we observed elevated expression of TACI on long-lived *ex vivo* plasma cells relative to short-lived ones **(Fig. S6 B)**. Though primary plasma cells express multiple TLR family members, 5TGM1 cells only expressed TLR4 **(Fig. S6 C)**. We therefore ablated *Tnfrsf13b*, *Tlr4*, *Myd88*, and *Traf6* (the gene that encodes TRAF6, a downstream molecule of the MYD88 cascade) in 5TGM1 myeloma cells using CRISPR-Cas9. Ablation of each of these genes resulted in reduced surface CD98 and fewer antibodies in culture supernatants relative to cells transduced with a control gRNA (**Fig. 9 A-B)**. To test if signaling through TACI and TLR ligands promote antibody secretion in *ex vivo* plasma cells, we treated purified short-and long-lived plasma cells from B6 mice with the High Mobility Group Box 1 (HMGB1) protein, a known endogenous ligand of TLRs (Yu et al., 2010), or B-cell Activating Factor (BAFF) or A Proliferation-inducing ligand (APRIL), known ligands of TACI, and measured secretory capacity in ELISpot assays. In parallel, we treated plasma cell subsets from *Myd88*-KO mice with the aforementioned ligands. Exogenous HMGB1 did not impact antibody secretion in any of the plasma cell subsets examined (**Fig. 9 C)**. Addition of BAFF or APRIL, however, increased the spot size of longer-lived plasma cells relative to untreated groups and had minimal impact on short-lived plasma cells (**Fig. 9 D-E)**. This effect on antibody secretion was MYD88-dependent, as these cytokines had no effect on plasma cells from *Myd88*-KO mice. Thus, APRIL and BAFF signal through TACI and MYD88 to influence the antibody secretory capacity of long-lived plasma cells.

**Figure 9:**
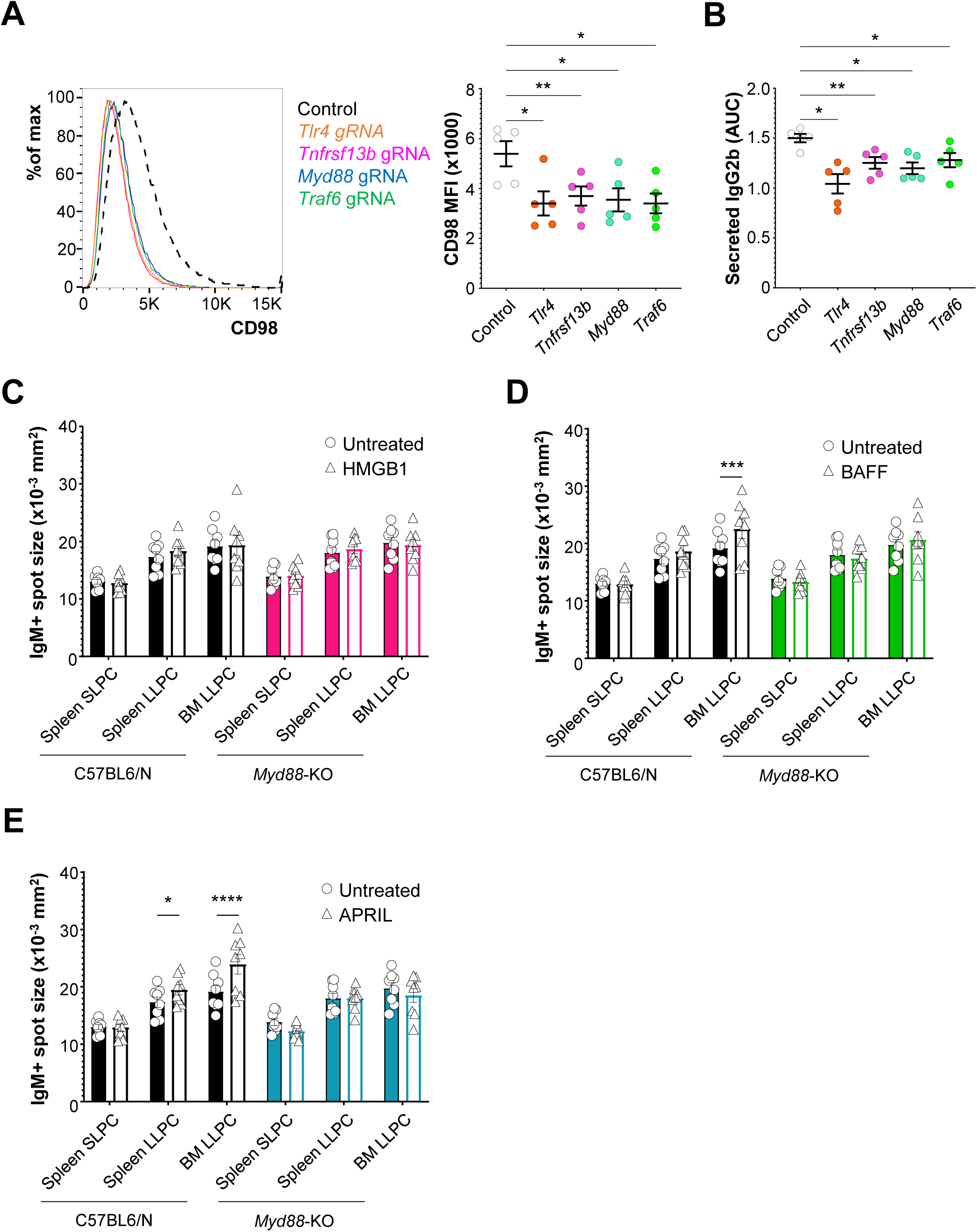
BAFF and APRIL promotes antibody secretory capacity in a MYD88-dependent manner. **(A)** 5TGM1-Cas9 cells were transduced with gRNA targeting *Tlr4*, *Tnfrsf13b*, *Myd88*, or *Traf6*. Representative histograms (left) showing surface CD98 levels in transduced groups (colored) relative to control gRNA transduced cells (black dashed). CD98 MFI (right) was quantified across 5 independent experiments. Each circle represents values from cells in a single well. *p<0.05 by repeated measures one-way ANOVA with Dunnett’s correction. **(B)** Secreted IgG2b from culture supernatant of groups in (A) were quantified by ELISA. AUC for each group is shown. Pooled data from 5 experiments. *p<0.05 by repeated measures one-way ANOVA with Dunnett’s correction. **(C-E)** Sorted plasma cell subsets from C57BL/6 or *Myd88*-KO mice were seeded into an ELISpot plate with media containing (C) 10µg/mL HMGB-1, (D) 10ng/mL BAFF, or (E) 10ng/mL APRIL. Cells treated with media are indicated with black bars, while treated groups are colored. Individual circles and triangles represent the mean IgM+ spot size from one mouse and matching groups are joined by a line. Pooled data from 8 mice across 4 experiments. *p<0.05 by Two-way ANOVA.

## Discussion

Plasma cell subsets vary in their lifespans and presumably their ability to sustain durable humoral immunity. Subtle differences in CD98 expression distinguish these subsets, thereby providing a convenient handle for unbiased genome-wide screens for factors that promote plasma cell longevity. We identified 34 genes that included a variety of vesicular proteins, subunits of the V-type ATPase complex, RNA export and processing proteins, and various signaling factors that promote CD98 expression. Though the original intent of the screen was to identify determinants of lifespan, all of the factors we tested impacted secretory capacity. These data suggest some convergence in factors that fine tune plasma cell lifespan and antibody secretion.

Nearly half of the genes identified in our screen encode proteins associated with intracellular vesicles and acidification. The vesicular network is relevant to plasma cells to support the secretory capacity and high rate of translation, folding, and antibody production in these cells (Casey et al., 2010). The frequency of acidic vesicles was higher in longer-lived plasma cell subsets relative to their short-lived counterparts, consistent with their elevated antibody secretory capacity. Impeding RAB7 exchange with RAB5 on endosomes inhibited antibody secretion and plasma cell longevity, demonstrating the importance of endosomal function in plasma cells (Pone et al., 2015; Lam et al., 2016). The greater numbers of acidic vesicles in long-lived plasma cells compared to short-lived plasma cells likely depend on the activity of the inositol kinase PI4KB, which has been shown in multiple systems to promote lysosomal integrity, vesicle formation, and retention of integral proteins to these organelles like the V-type ATPase complexes (Balla et al., 2002; Sridhar et al., 2013). The factors that might promote differential PI4KB activity in plasma cell subsets are currently unknown.

Our data further reveals the complex regulation of antibody secretion in plasma cells by identifying a role for the RNA helicase DDX3X in modulating this process. This protein has been shown to associate with secondary structures at the 5’ end of mRNA and unwinds them through its helicase activity, allowing for recruitment of ribosomes and its subsequent translation (Calviello et al., 2021; Shih et al., 2008). In plasma cells, this function is particularly relevant as DDX3X activity might ease barriers in mRNA translation and support a steady rate of antibody synthesis and secretion. While it is likely that DDX3X associates with *Igh* and *Igl* mRNA due to their sheer abundance in plasma cells (Shi et al., 2015), identifying transcripts bound to DDX3X in plasma cells might provide insights to the translational regulation of key proteins for cellular function. As mice with lymphocyte-specific ablation of *Ddx3x* show reductions in B cell numbers (Szappanos et al., 2018; Liu et al., 2018), generating inducible models of *Ddx3x* deletion may help us understand the role of this factor in plasma cell longevity. Further, as males express a homolog of DDX3X called DDX3Y, it would be interesting to observe if the two factors have overlapping targets in plasma cells from males and compensate for each other in the absence of one or the other (Sekiguchi et al., 2004).

While MYD88 has been well-studied in B cells, our report demonstrates a plasma cell intrinsic role for this signaling intermediate in modulation CD98 expression, survival, and antibody secretion. We further demonstrate that BAFF and APRIL likely mediate these effects through TACI and MYD88, but not through interactions with their other receptors, BAFF-R and BCMA, both of which are MYD88-independent (He et al., 2010; Smulski and Eibel, 2018). Signals through APRIL and its receptors are particularly relevant in the bone marrow niche, where plasma cells are immersed in a milieu of extracellular matrix proteins, cytokines, and cell adhesion factors that promote their survival *in vivo* (Cassese et al., 2003; O’Connor et al., 2004; Belnoue et al., 2008; Rozanski et al., 2011; Nguyen et al., 2018; Ishikawa et al., 2024). BAFF in turn has been shown to be crucial for B cell survival and plasma cell differentiation (Avery et al., 2003; Benson et al., 2008; Parsa et al., 2016). Our findings suggest that while APRIL can promote plasma cell survival by interacting with both BCMA and TACI, and BAFF with TACI and BAFF-R (Eslami et al., 2024), they can only enhance antibody secretion through their interaction with TACI and subsequent signaling through MYD88. This data also indicates that the secretory capacity of plasma cells is not exclusively a cell intrinsic function but can be modulated by external stimuli.

CRISPR-Cas9 screens were recently carried out by multiple independent groups to identify factors regulating plasma cell differentiation (Pinter et al., 2022; Xiong et al., 2022) and antibody secretion (Trezise et al., 2023) using *in vitro* primary B cell systems. Of the genes shown to be important for antibody secretion, such as those involved in the unfolded protein response, we found little overlap with the genes in our dataset. This may be due to our use of CD98 as a marker, which varies with antibody secretory capacity across plasma cell subsets but not as a function of the unfolded protein response (Lam et al., 2018). Our screen was thus more likely to identify pathways that differ between plasma cell subsets than common pathways required for secretory capacity. Put together, our data suggest there are distinct programs required for the induction of antibody secretion: those that are absolutely required and used similarly across plasma cell subsets, and others that further tune lifespan and secretory capacity. At least some of the latter pathways functionally link survival and antibody secretion, thereby allowing relatively few highly secretory long-lived plasma cells to provide robust protection.

## Acknowledgements

The authors wish to thank the flow cytometry core at the University of Arizona for their assistance. The core is supported by RII of the University of Arizona and an NCI grant P30CA023074. Use of the Imagestream was made possible by a NIH award S10 OD028466. We are also grateful to Colin Fields for his assistance in procuring human samples after surgery and Jean Wilson, Hannah Pizzato, and Kit Tobey for their inputs. We also wish to thank William Smith and the staff at University Animal Care for their assistance with mouse husbandry and maintenance. This project was supported by a NIH grant R01AI129945 to D.B. and a Bio5 postdoctoral fellowship to L.J.D. The funders of this project had no role in study design, data collection and interpretation, decision to publish, or composing the manuscript. There was no additional external funding received for this study.

## Materials and methods

### Mice

C57BL/6N mice (556) were purchased from the Charles River laboratories. *Myd88*-null (9088) and *Myd88*-flox (8888) mice were purchased from Jackson Laboratories (Hou et al., 2008). Jchain-CreERT2 mice have been generated previously (Wong et al., 2020). Mice were housed and bred under specific pathogen free conditions. Experiments were carried out on age- and sex-matched mice between 8-12 weeks in age. All animal procedures were executed under the guidelines provided by the Institutional Animal Ethics committee of the University of Arizona. Euthanasia was carried out by carbon dioxide asphyxiation at the rate of 1.8-4.0 L/minute in a 7L chamber until 1 minute after respiration ceased. Mice were then cervically dislocated to ensure death. For some experiments, mice sedated with Fluriso (Vet One) were injected intravenously with 100μg of 2NBDG (Cayman Chemical company) and euthanized after 20 minutes.

### Human tissues

Tonsil samples were obtained from consenting adults undergoing elective tonsillectomies (Banner-University Medical Center). Bone marrow samples were obtained from persons undergoing robotics-assisted hip arthroplasty (Tucson Orthopedic Institute). All donors were anonymous, and no patient data was collected as part of this investigation. All human work was conducted in accordance with federal, state, and county regulations and approved by the University of Arizona IRB.

### Immunizations

Mice were immunized intraperitoneally with 100µg NP-CGG (Biosearch Technologies) in 1% Alhydrogel (Invivogen). Circulating blood was collected by venipuncture of the tail vein from immunized mice and allowed to stand at room temperature to coagulate overnight. Serum was collected after spinning down the samples at 15000rpm for 10 minutes. At 8 weeks post-immunization, mice were fed special chow containing 400mg Tamoxifen citrate per kg diet (Envigo) for two weeks prior to euthanasia.

### Primary cells

Single cell suspensions of spleens were prepared by macerating organs with frosted slides while bone marrow cells were isolated from tibiae, fibulae, femurs, and pelvic bones using a mortar and pestle in 1x PBS containing 5% Adult bovine serum (MP Biomedical; hereon referred to as FACS buffer). Human tonsils were sliced using a scalpel and forceps and ground in FACS buffer using a mortar and pestle. Human bone marrow reamings were shaken vigorously in a sterile container containing FACS buffer. In all cases, cells were filtered through a 70-micron nylon mesh to remove debris. Single cell suspensions were treated in an ammonium chloride-potassium (ACK) hypotonic lysis solution to lyse erythrocytes followed by density gradient centrifugation on a Histopaque-1119 layer (Millipore Sigma). To enrich for murine plasma cells, whole cell suspensions were enriched by labelling with an appropriate anti-CD138 antibody (Biolegend) followed by anti-APC or anti-PE microbeads and enriched using LS columns (all from Miltenyi Biotec). CD38+ tonsil and bone marrow cells were enriched similarly from whole cell suspensions. For some experiments, CD38-enriched cells were cryopreserved in 50% incomplete RPMI (Thermo Fisher Scientific), 40% fetal calf serum (Peak Serum), and 10% DMSO (Millipore Sigma) until the time of experimentation. For intracellular staining, cells were first fixed with 2% Paraformaldehyde (Electron microscopy services) for 10 minutes followed by permeabilization with 0.1% Saponin (Millipore Sigma).

### Plasmids

The mouse Brie CRISPR-knockout library (73633), lentiGuide-puro (52963), pLenti-sfGFP-LAMP1-mCherry (164478), and pHAGE-DDX3X (116730) were all purchased from Addgene (Doench et al., 2016; Sanjana et al., 2014; Webb et al., 2021; Ng et al., 2018). The lentiGuide-mCherry plasmid (217005) was generated in-house (D’Souza et al., 2022). Edit-R inducible lentiviral hEF1α-Blast-Cas9 nuclease plasmid (CAS11229) was purchased from Dharmacon. Guide RNA (gRNA) targeting genes of interest were chosen from the Brie CRISPR-knockout library and cloned into an empty lentiGuide-puro construct as described previously (D’Souza et al., 2022). Mouse genomic *Ddx3x*-specific gRNA were designed using CRISPick (Doench et al., 2016; Sanson et al., 2018). pMD2.G (12259) and psPAX2 (12260) were used to generate lentivirus and were also purchased from Addgene. T323I, R326H, I415Δ, and T532M mutations were introduced into the parental pHAGE-DDX3X by PCR and site-directed mutagenesis using the KLD enzyme reaction (New England Biolabs). Positive mutants were identified by Sanger sequencing (Eton Biosciences). Plasmids were transformed and grown in XL-1 Blue cells (Agilent Technologies) or *Stbl4* competent cells (Thermo Fisher Scientific) for plasmids larger than 10kb.

### Cell lines and cell culture

The 5TGM1 myeloma line was a kind gift from Michael H. Tomasson while at Washington University in St. Louis (Garrett et al., 1997). 5TGM1 cells expressing a Doxycyline-inducible Cas9 (5TGM1-iCas9) was generated by transducing cells with lentivirus containing the Edit-R inducible Cas9 construct at 2500rpm for 90 minutes at room temperature followed by culturing cells in media containing 10µg/mL Blasticidin-HCl (Thermo Fisher Scientific). Cells were sorted into single cell clones by FACS and screened for lines not expressing Cas9 in regular doxycycline-free media. 5TGM1 cells expressing Cas9 constitutively (5TGM1-Cas9) were generated previously (D’Souza et al., 2022). These cells were transduced with lentivirus containing the pHLARE cassette by spin-infection and positive cells were FACS-sorted as GFP+ mCherry+ cells. Overexpression constructs for human DDX3X and mutants were introduced into 5TGM1-Cas9 cells similarly and purified based on GFP expression. All 5TGM1 lines were cultured in RPMI (Gibco) containing 10% FBS, 1mM Sodium pyruvate, 2mM GlutaMAX, 10µg/mL Ciprofloxacin-HCl, minimum-essential amino acids, Penicillin and Streptomycin (hereon referred to as cRPMI). After introduction of gRNA constructs, cells were incubated for 5 days at 37°C with 5% CO_2_. For antibody secretion assays, 100,000 5TGM1 cells were seeded in triplicates into wells of a 96-well plated and incubated overnight. Culture supernatants were harvested the next day and assayed for secreted immunoglobulins. Where indicated, cells were treated with Bafilomycin A1 (Millipore Sigma) at varying concentrations. For some experiments, cells were treated with 10µg/mL mouse HMGB-1 (ACRO Biosystems), 10ng/mL recombinant mouse BAFF (R&D Systems), or 10ng/mL recombinant mouse APRIL (Thermo Fisher Scientific). Lenti-X 293T cells were purchased from Takara and cultured in DMEM (Cytiva) with all the additional supplements described previously for RPMI. Lentivirus were generated from these cells by transient transfection of plasmids with packaging constructs previously complexed with GeneJuice (Millipore Sigma).

### Flow cytometry

Single cell preparations were stained with the following antibodies from Biolegend: anti-mouse CD138-PE, -APC, or -BV510 (281-2); anti-mouse B220-AlexaFluor700 (RA3-6B2); anti-mouse CD98-AlexaFluor647 or -PE-Cy7 (4F2); anti-human CD38-PE (HB-7); anti-human CD27-APC (M-T271); anti-human CD19-BV421 (HIB19); anti-human IgM-BV650 (MHM-88); anti-human CD138-FITC (MI15). Rat anti-mouse IgA-BV421 (C10-1) was purchased from BD Biosciences. Goat F(ab’)2 anti-human IgA-AlexaFluor647 (2052-31) was purchased from Southern Biotech. Unconjugated antibodies against mouse/human ATP6V1A (EPR19270), RAB7 (EPR7589), and LAMP1 (ab24170) were all purchased from Abcam and used at a 1:500 dilution. Stained organelles were detected with an anti-rabbit IgG-AlexaFluor405 reagent (A48258, Thermo Fisher Scientific). Other antibodies from Thermo Fisher Scientific used in this study are anti-mouse IgM-PerCP-eFluor710 (II41), and anti-mouse/human DDX3X (A300-474A). Propidium iodide and DAPI (both from Millipore Sigma), were used to stain dead cells in some assays. In some experiments Zombie UV or Zombie Red (both from Biolegend) were used to stain dead cells prior to fixation and permeabilization. Cells were analyzed for fluorescence on a BD Fortessa cytometer (BD Biosciences) and data interpreted using the FlowJo software (BD Biosciences). Intracellular organelle frequencies in cells were examined on an Imagestream^X^ MkII (Cytek Biosciences) at 60x magnification with extended depth of field. Spot numbers per cell and similarity morphology indexes were calculated on the IDEAS software (Cytek Biosciences). Fluorescence associated cell sorting was carried out on a FACS Aria II (BD Biosciences).

### Library screening and next generation sequencing

The mouse Brie library was packaged into lentiviral particles by transient transfection of Lenti-X 293T cells with the 5µg of the library plasmid, 1.75µg of pMD2.G and 3.25µg of psPAX2 packaging constructs complexed with the GeneJuice transfection reagent (Millipore Sigma). Culture supernatants were collected and pooled at 48h and 96h and filtered through a 0.45µ syringe filter. Fresh filtered supernatants containing virus were then used to transduce 5TGM1-iCas9 cells in the presence of 8µg of Polybrene (Millipore Sigma). Positively transduced cells were then selected in cRPMI containing Puromycin (Gibco). Cas9 expression in 40 million positively transduced 5TGM1-iCas9 cells (roughly 500x coverage of plasmid library) was induced after adding 300ng/mL Doxycycline hyclate to culture media. After 7 days, cells were cultured with 10µg/mL of 2NBDG and then stained for surface CD98 expression. Live CD138+ CD98-low cells were then FACS sorted into individual 15mL conical tubes. Genomic DNA from cells were extracted using the mini genomic DNA extraction kit (IBI scientific) and gRNA constructs amplified from it using Q5 Polymerase (New England Biolabs). Flow cell attachment sequences, Illumina sequencing regions, and unique 3’ barcode sequences were then introduced by PCR. These pair-ended 150bp libraries were then sequenced on a Novaseq 6000 (Illumina). FASTQ files containing 325-926 million reads were then analyzed for gene enrichment in sorted fractions using the Python-based MaGeCK module (Li et al., 2014). Representation of the Brie library in cells were confirmed by comparing representation of gRNA from positively transduced cells to the parental Brie library. Volcano plots showing enriched genes in sorted fractions relative to unenriched cells were plotted using Prism v.10 (Graphpad).

### ELISA

ELISA plates (Corning) were coated overnight at 4°C with unconjugated rat anti-mouse κ-light chain (187.1, BD Biosciences) in a 0.1M Sodium carbonate-bicarbonate buffer (pH: 9.5). The next day, plates were blocked with 2% Bovine Serum Albumin (Millipore Sigma) in 1xPBS containing 0.05% Tween-20 (1xPBS-T) followed by incubation with serial dilutions of cell-free culture supernatants. After 3 hours, plates were then washed thrice with 1xPBS-T followed by an overnight incubation at 4°C with biotinylated rat anti-mouse IgG2b (SB74g, Southern Biotech). Plates were washed the next day in 1xPBS-T followed by an incubation with Streptavidin linked to horseradish peroxidase (BD Biosciences) for an hour at room temperature. Plates were developed using a TMB substrate (Alfa Aesar) and arrested with 2N Sulfuric acid (Millipore Sigma). Absorbance was measured at 450nm and corrected for background absorbance at 650nm on a VERSAmax microplate reader (Molecular devices). Area under the curve (AUC) was calculated from background adjusted A450 values on the Prism v.10.3 (Graphpad). A similar workflow was used for detection of NP-specific antibodies in serum, except that ELISA plates were coated with NP_10-19_-BSA (Biosearch technologies) and detected with a Peroxidase-conjugated donkey anti-mouse IgG(H+L) (Jackson ImmunoResearch).

### ELISpot

Multiscreen HTS HA filter plates (Millipore Sigma) were coated with NP-BSA (Biosearch technologies), Goat anti-mouse Ig(H+L) (Southern Biotech), or a 1:1 mix of Goat anti-human Lambda and Kappa (both from Southern Biotech) prepared in 1xPBS. After an overnight incubation at 4°C, plates were blocked with RPMI containing 10% FBS at 37°C. Sorted cells were resuspended in cRPMI and seeded in triplicates in their designated wells in the plate and incubated at 37°C overnight. Plates were washed the next day with 1xPBS and then incubated overnight with the following peroxidase-conjugated secondary reagents as appropriate: donkey anti-mouse IgG(H+L) for NP-specific plasma cells (Jackson Immunoresearch); goat anti-mouse IgM (1020-05), anti-IgG (1030-05), and anti-IgA (1040-05) for mouse splenic and bone marrow plasma cells; goat anti-human IgM (2020-05), anti-IgG (2045-05), and anti-IgA (2050-05) for detection of human bone marrow plasma cells (all from Southern Biotech). Spots were developed using a TrueBlue Peroxidase substrate (Kirkegaard and Perry Laboratories) and washed vigorously with MilliQ water. Spot size and numbers were quantified on the CTL ImmunoSpot S6 analyzer. The total numbers of NP+ plasma cells were calculated by multiplying the number of spots with the dilution factor and divided by total bone marrow cell counts.

### Statistical analysis

All statistical analysis was done using Prism v. 10.3 (Graphpad). Specific statistical tests and significance are indicated in the figure legends of the indicated graphs. Fold changes and p-values for the genome-wide screen were calculated by MaGeCK (Li et al., 2014). Figure S2 A was generated using Biorender.

---

**Supplementary Figure 1:**
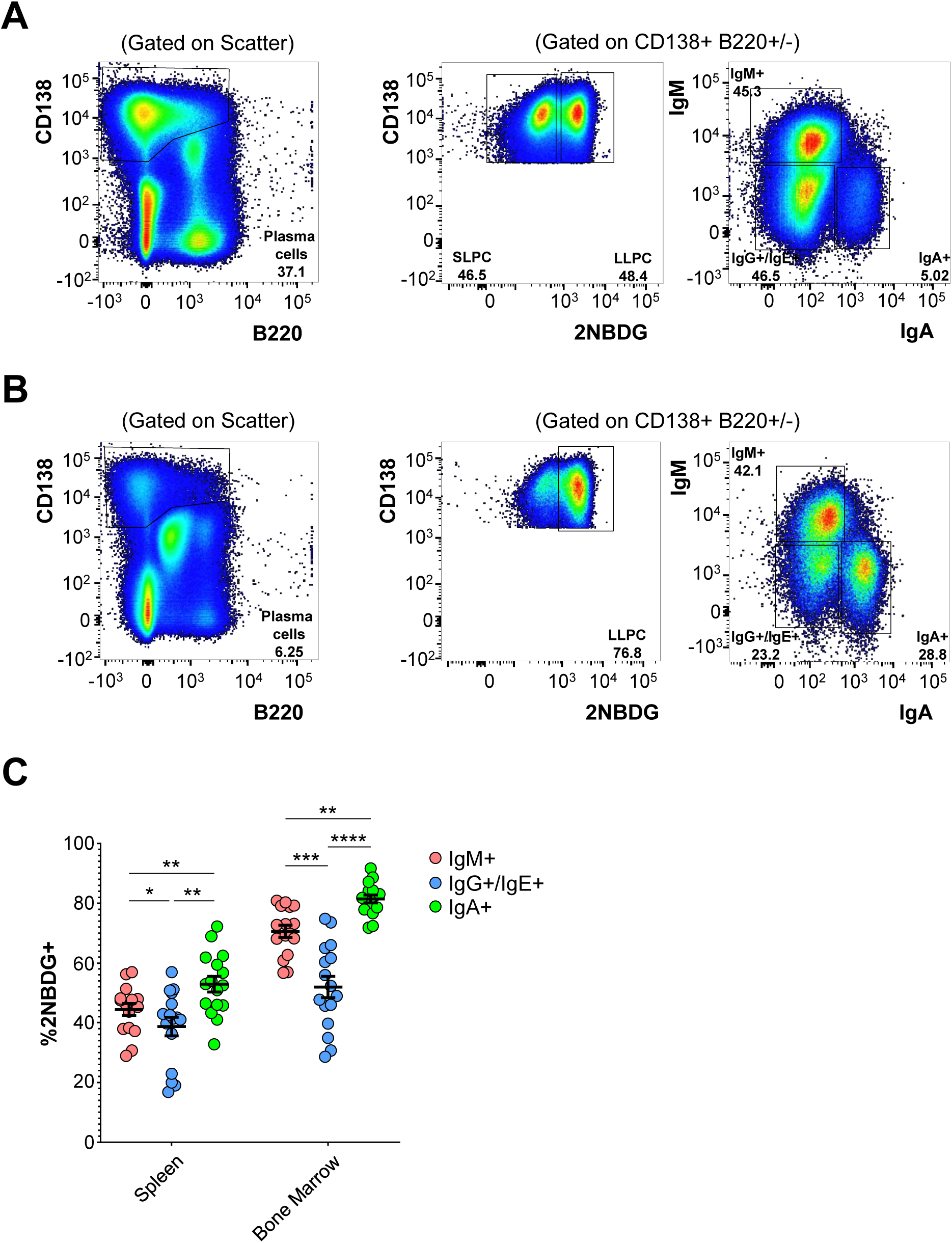
Murine plasma cell subsets have disparate proportions of 2NBDG+ cells. **(A)** Representative gating of mouse splenic plasma cells on enriched CD138+ splenocytes. Single cells were gated first as CD138+ B220+/-cells (left) and then distinguished as 2NBDG-SLPC and 2NBDG+ LLPC (center), IgM+, IgA+, or IgG+ and IgE+ (right). **(B)** Representative gating of murine bone marrow plasma cell subsets gated as in (A). **(C)** Long lived plasma cell frequencies as defined by 2NBDG uptake in mouse splenic and bone marrow IgM+, IgG+ and IgE+, and IgA+ plasma cells. Each circle represents values for the respective subset from one mouse. Pooled data from 16 mice across 4 experiments. *p<0.05 by two-way ANOVA with Tukey’s correction.

**Supplementary Figure 2:**
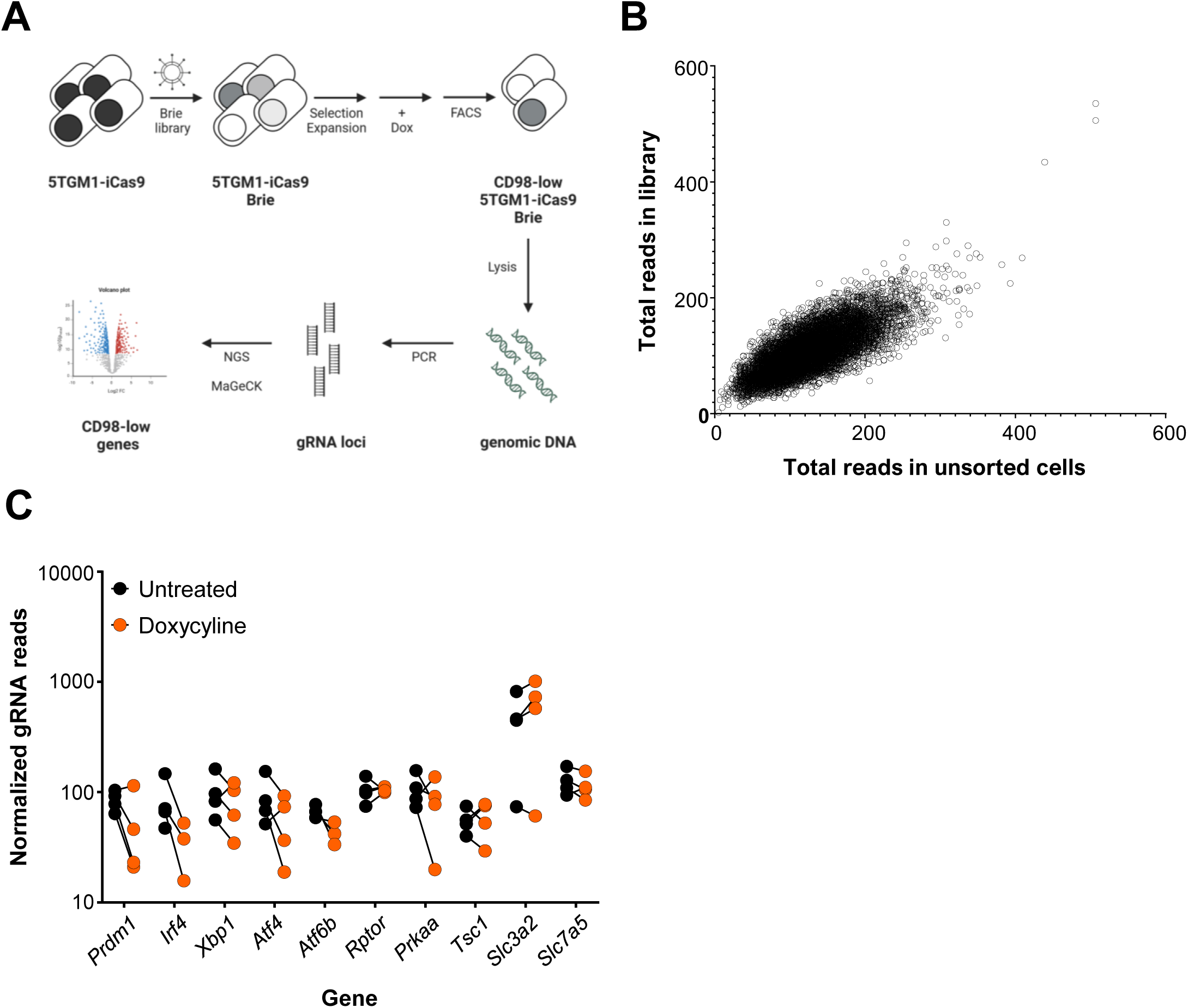
Cas9 induction leads to reduced abundance of gRNAs targeting essential genes in 5TGM1-iCas9-Brie cultures. **(A)** Schematic of genome-wide unbiased CRISPR-Cas9 screen in 5TGM1 cells. **(B)** Scatter plot of gRNA read counts following next-generation sequencing of the mouse Brie library (Y axis) against the corresponding read numbers in uninduced 5TGM1-iCas9-Brie cells (X axis). Each open circle represents one gRNA. **(C)** 5TGM1-iCas9-Brie cells were left untreated or treated with Doxycycline. At day 7, gRNA abundance was quantified from both groups using MaGeCK. Normalized counts of gRNA targeting indicated genes are shown in both groups. Matching gRNAs are connected by a line. Pooled data from 4 biological replicates.

**Supplementary Figure 3:**
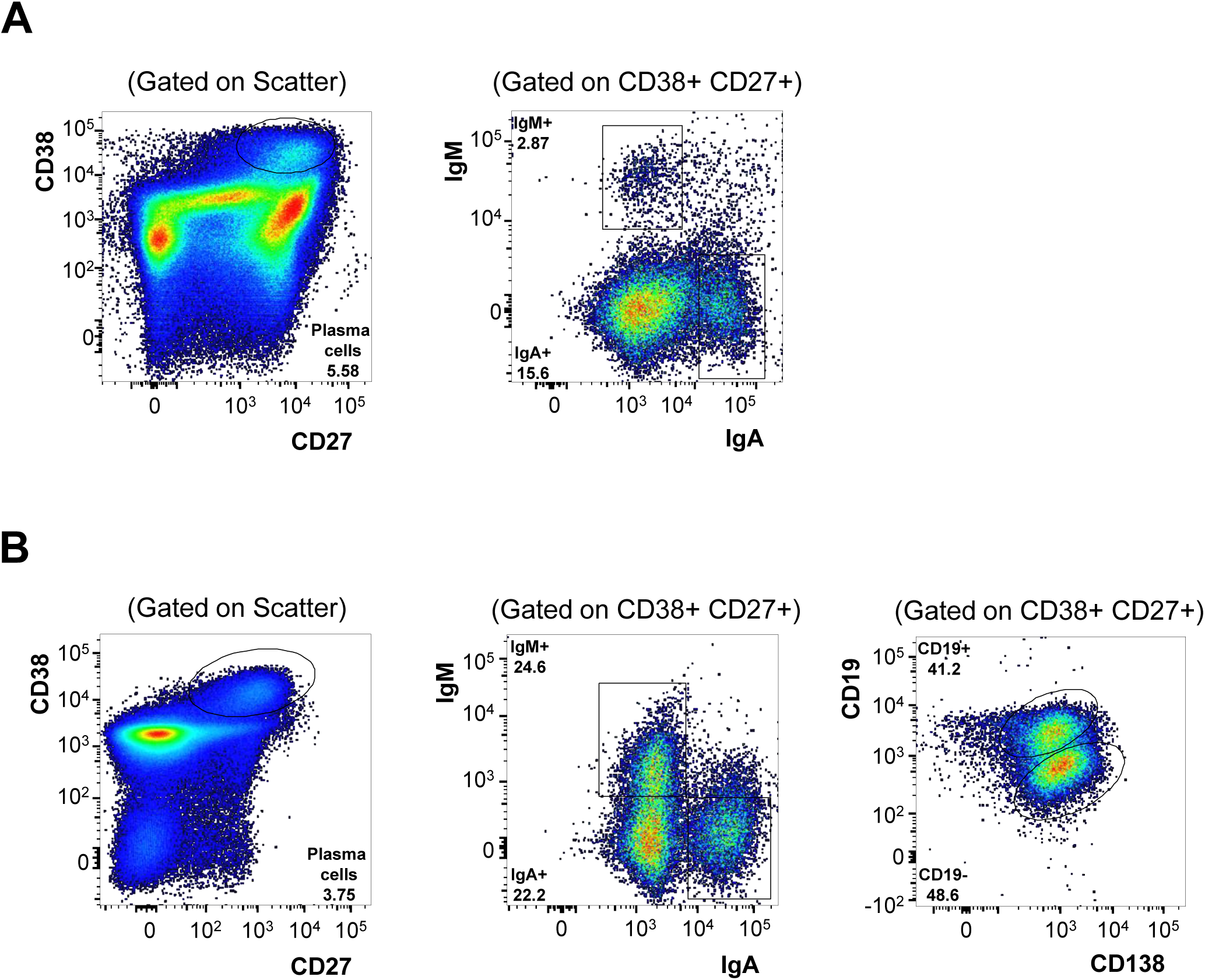
Identification of human plasma cell subsets. **(A)** Representative gating of human tonsil plasma cells on CD38+ enriched cells. Cells were identified as CD38+ CD27+ cells on single cells (left). Cells were then identified as IgM- or IgA-secretors based surface expression of the respective isotype (right). Double negative cells were presumed IgG+ and IgE+ plasma cells. **(B)** Bone marrow plasma cells of various isotypes were gated like tonsillar plasma cells as CD38+ CD27+ single cells (left, center). Human plasma cell subsets were then identified as CD19+ or CD19-cells respectively (right).

**Supplementary Figure 4:**
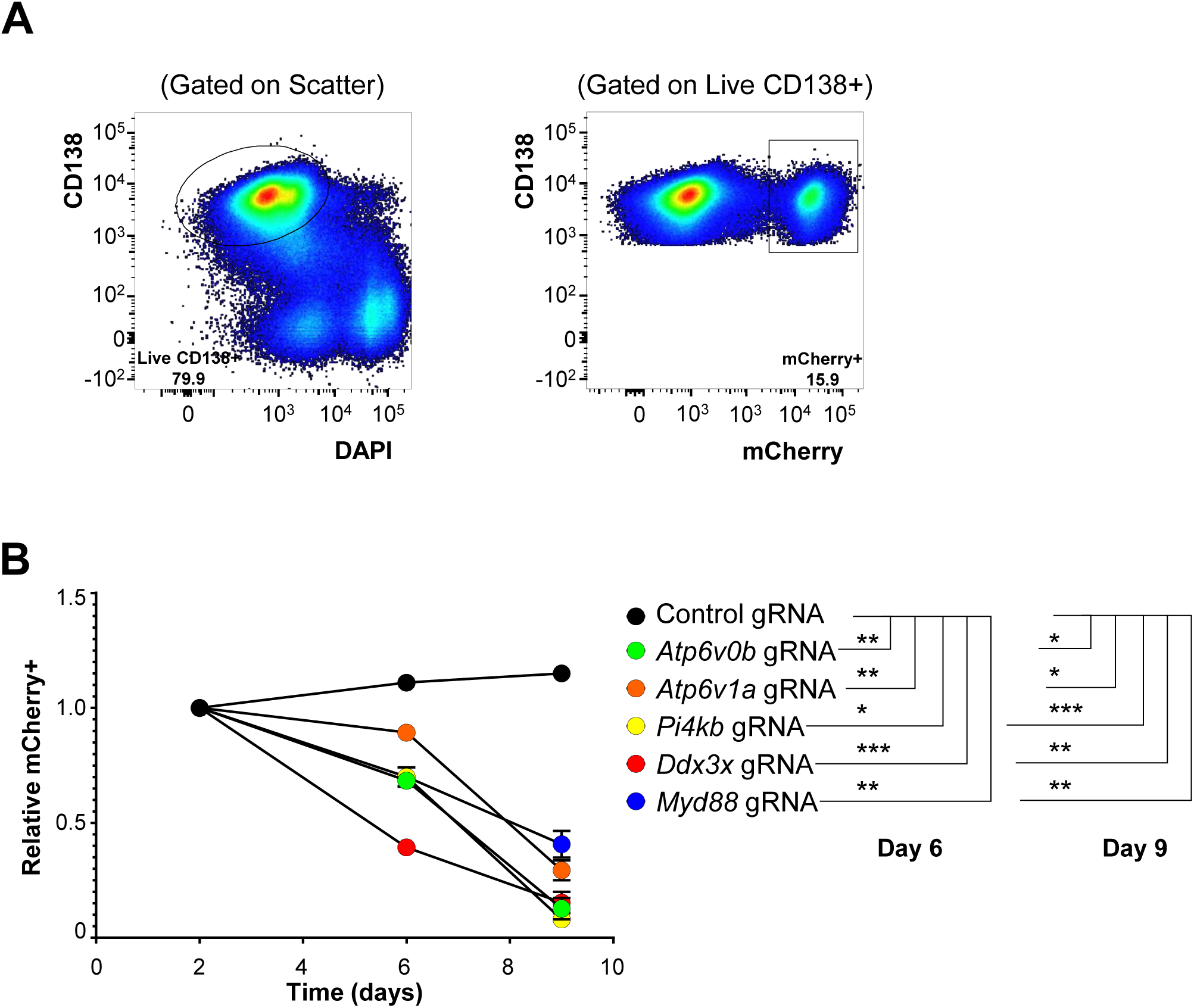
Perturbations in *Atp6v0b*, *Atp6v1a*, *Pi4kb*, *Ddx3x*, and *Myd88* compromise cell survival. **(A)** 5TGM1-Cas9 cells were transduced with lentivirus containing lentiGuide-mCherry with control gRNA or gRNA targeting *Atp6v0b*, *Atp6v1a*, *Pi4kb*, *Ddx3x*, and *Myd88*. Representative flow cytometry plots depicting live myeloma cells (left) at day 2 post transduction with mCherry+ frequencies on them (right). **(B)** Transduced cells in (A) were tracked in culture over time and mCherry+ frequencies measured at days 2, 6, and 9 after transduction. mCherry+ frequencies were normalized to frequencies at day 2. Each circle represents mCherry frequencies of cells in that well and groups across time are linked by a line. Statistical significance for each group relative to the control was calculated and indicated in the figure legend for day 6 and day 9. *p<0.05 by two-way ANOVA.

**Supplementary Figure 5:**
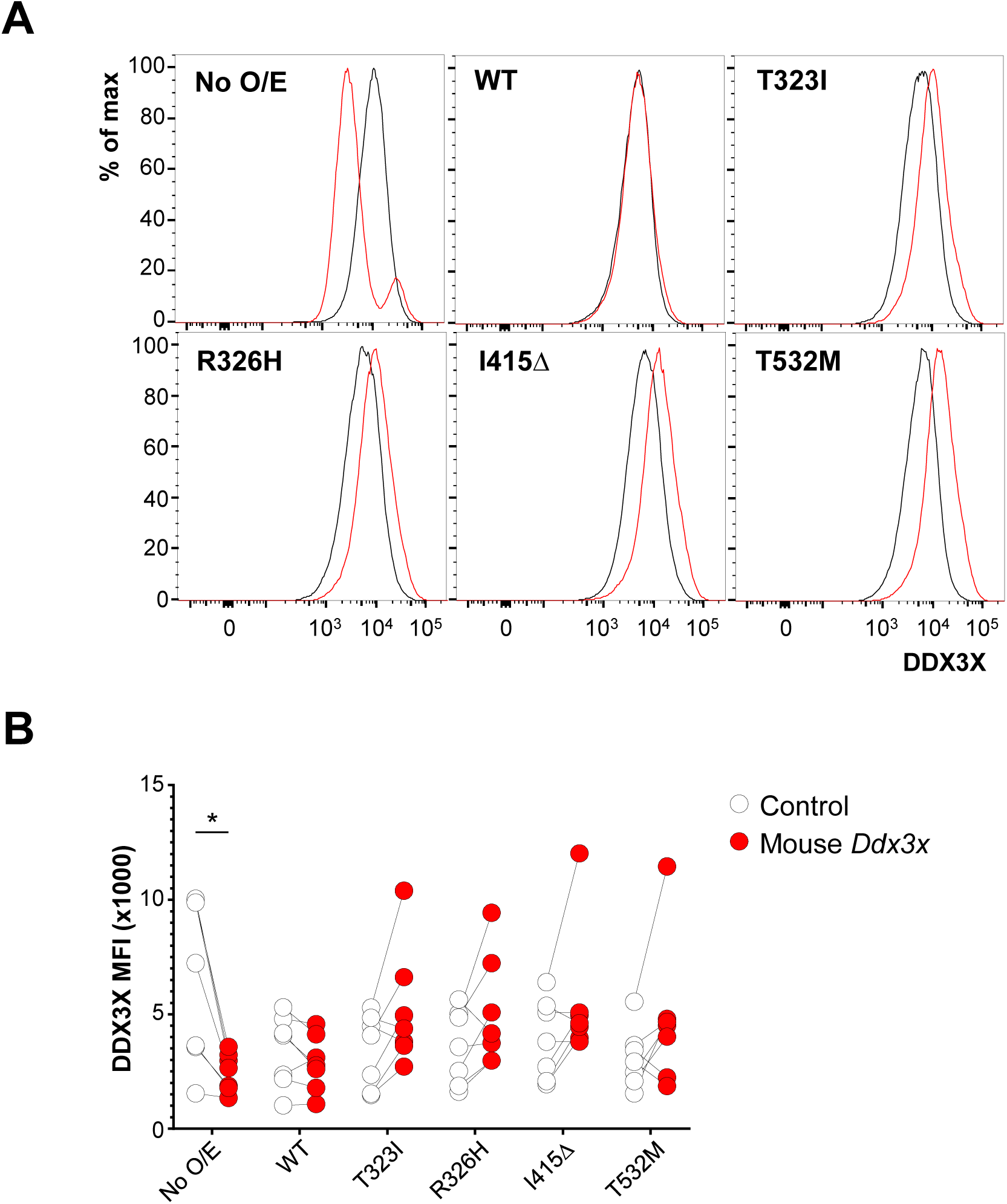
DDX3X protein levels are comparable in overexpression cultures after ablation of endogenous mouse DDX3X. **(A)** 5TGM1-Cas9 cells overexpressing human DDX3X proteins were transduced with gRNA targeting mouse *Ddx3x* (red) or control gRNA (black). Cells not overexpressing DDX3X are shown on the top left. Representative histograms are shown for each of the groups. **(B)** Quantification of the MFI of various groups in (A) are displayed. Each line connects groups within the same experiment. Pooled data from 7 independent experiments. *p<0.05 by two-way ANOVA with Sidak’s correction.

**Supplementary Figure 6:**
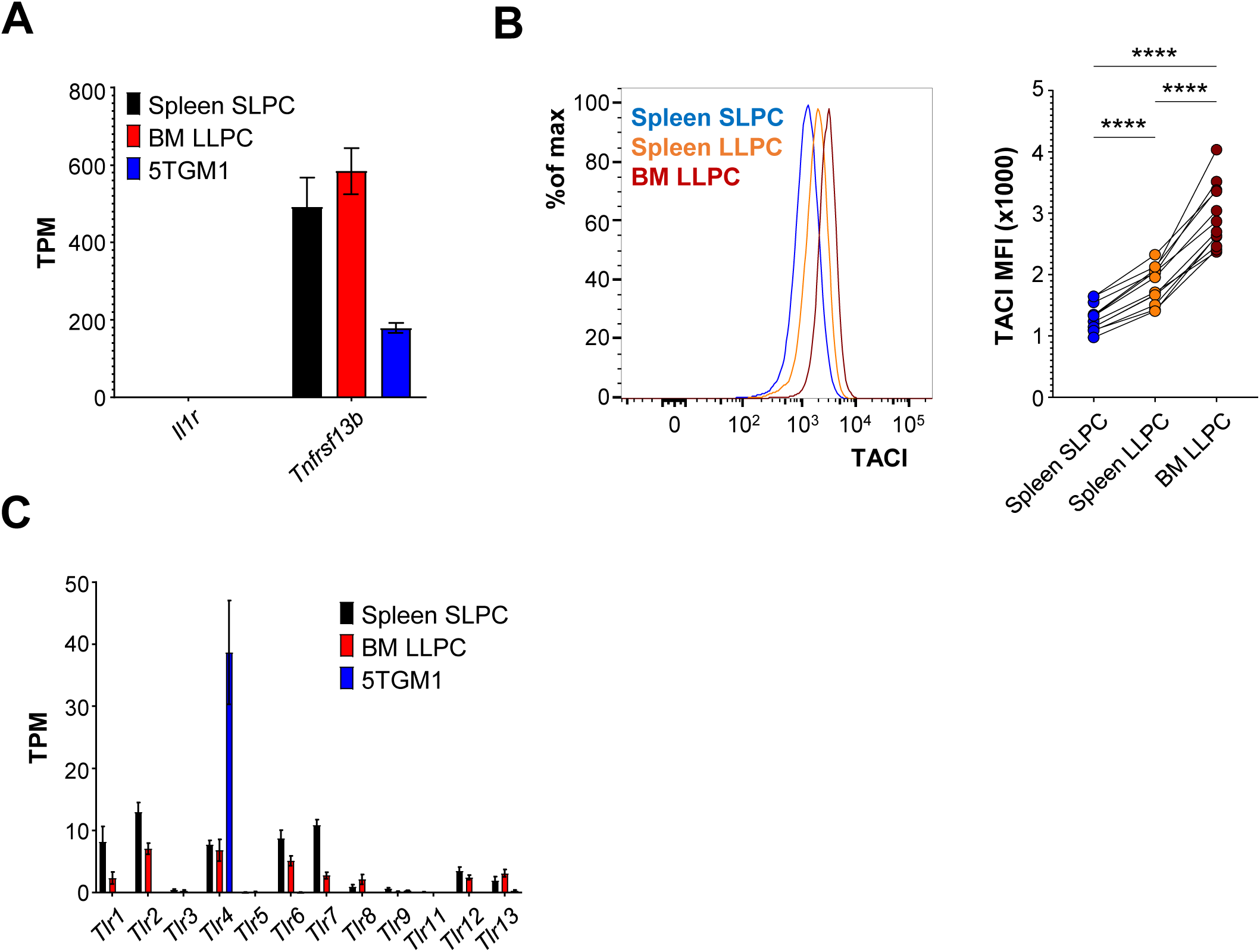
Plasma cell subsets express TACI and multiple TLRs but not IL-1R. **(A)** TPM values of IL-1R (*Il1ra*) and TACI (*Tnfrsf13b*) showing data from splenic and bone marrow plasma cell subsets from Lam WY *et al* (Cell Reports 2018) and 5TGM1 cells from D’Souza L *et al*, (PLoS One 2022). Pooled data from three independent experiments. **(B)** Representative histogram (left) showing surface expression of TACI on murine plasma cell subsets as measured by flow cytometry. Each data point represents cells from one mouse, and subsets from each mouse are connected by lines. Pooled data from 12 mice across 7 experiments. *p<0.05 by paired one-way ANOVA with Tukey’s correction. **(C)** TPM values of members of the TLR family of genes are shown for populations mentioned in (A). Pooled data from three independent experiments.

